# Closed-loop acoustic stimulation promotes local slow oscillation train initiation and persistence in elderly adults

**DOI:** 10.64898/2026.10.29.762885

**Authors:** Lucila Capurro, Leonela M. Tassone, Michael Radloff, Luis I. Brusco, Rodrigo Ramele, Cecilia Forcato

## Abstract

Slow oscillations (SOs) during non-rapid eye movement (NREM) sleep support memory consolidation and metabolic waste clearance. Aging is associated with reduced SO amplitude and density and disrupted temporal organization, characterized by more isolated events and shorter SO trains. We investigated whether closed-loop acoustic stimulation (CLAS) could enhance the initiation and persistence of rhythmic SO activity in elderly adults. Twenty-seven participants underwent a baseline night followed by an experimental night, during which online-detected SOs were either stimulated using CLAS (STIM; n = 16) or left unstimulated (SHAM; n = 11). We quantified the proportions of isolated and consecutive SOs, train-initiation probability, and SO train length. Analyses assessed local effects around online-detected SOs and global effects across NREM sleep using offline detection. Locally, CLAS reduced the proportion of isolated SOs and increased the proportion of consecutive SOs, train-initiation probability, and mean train length. Global overnight effects were weaker but directionally consistent, primarily reflecting a shift toward longer SO trains. These changes occurred without a significant between-group difference in the change in SO density, and local effects remained significant after adjustment for SO density. Thus, CLAS promoted sustained rhythmic activity beyond changes in SO occurrence. These findings suggest that CLAS can modify the temporal organization of SOs in elderly adults, supporting its potential to mitigate age-related disruptions in SO dynamics. Temporal organization measures may complement conventional amplitude and density measures when evaluating CLAS effects. Further studies should establish whether these changes translate into cognitive benefits or improvements in other aspects of brain health.

## Introduction

Sleep slow oscillations (SOs; 0.5-1 Hz), prominent during non-rapid eye movement (NREM) sleep, are large-amplitude cortical slow waves characterized by alternating up and down states of neuronal excitability (Steriade et al., 1993; Achermann & Borbély, 1997). These oscillations play a central role in coordinating neural activity during sleep and are implicated in many physiological functions including synaptic plasticity and homeostasis, memory consolidation, immune function and glymphatic clearance (Diekelmann & Born, 2010; Rasch & Born, 2013; Tononi & Cirelli, 2014; Besedovsky et al., 2012; Xie et al., 2013).

Aging is associated with substantial alterations in sleep architecture and NREM slow oscillations, including reductions in amplitude, density, power, and synchronization (Mander et al., 2017; Scullin & Bliwise, 2015; Helfrich et al., 2018). Beyond these well-established changes, recent findings from our group demonstrated that aging also disrupts the temporal organization of SOs. Specifically, elderly adults exhibit a higher prevalence of isolated SOs, fewer SO trains, and shorter train lengths compared to younger adults (Capurro et al., 2026). These findings suggest a reduced capacity of cortical networks to sustain rhythmic oscillatory activity across time.

Closed-loop acoustic stimulation (CLAS) has emerged as a non-invasive neuromodulatory approach aimed at enhancing SO activity during sleep (Ngo et al., 2013; Santostasi et al., 2016). In CLAS paradigms, ongoing electroencephalographic activity is continuously monitored to detect the occurrence and phase of spontaneous SOs in real time. Brief auditory stimuli, typically pink noise bursts, are then delivered at specific phases of the oscillation, most commonly during the transition toward or around the up-state, where cortical excitability is maximal and stimulation appears to be most effective in amplifying subsequent oscillatory activity (Ngo et al., 2013; Ong et al., 2018; Wunderlin et al., 2021).

Since the pioneering studies demonstrating that phase-targeted auditory stimulation can enhance SO activity and improve memory consolidation (Ngo et al., 2013), a variety of CLAS protocols have been developed. These include single-stimulus paradigms, in which one auditory pulse is delivered following the detection of a SO; paired-pulse protocols, where a second stimulus is presented after the evoked oscillatory response to the first pulse (Ngo et al., 2015); and more sophisticated adaptive approaches that continuously estimate oscillatory phase and adjust stimulation timing in real time (Santostasi et al., 2016; Debellemaniere et al., 2018). Protocol duration also varies substantially across studies, ranging from brief stimulation blocks interleaved with silent periods to interventions spanning large portions of NREM sleep or even multiple consecutive nights (Papalambros et al., 2017; Wunderlin et al., 2021).

Evidence for the electrophysiological effects of CLAS has been more consistent in healthy young adults, with studies reporting enhanced SO activity, increased spindle activity, and stronger temporal coordination between SOs and spindles (Ngo et al., 2013, 2015; Ong et al., 2016). Several studies have also reported improvements in sleep-dependent memory consolidation, although the magnitude and reproducibility of these behavioral benefits remain variable across stimulation protocols (Ngo et al., 2013; Henin et al., 2019).

In aging populations, findings have been more variable. Papalambros et al. (2017) reported enhanced slow-wave activity (0.5-4 Hz) and improved declarative memory retention in elderly adults, whereas Schneider et al. (2020) found attenuated SO and spindle responses relative to young adults and no memory benefits in a middle-aged cohort. These findings suggest that the magnitude and functional consequences of CLAS effects may depend on both the population studied and the stimulation protocol, limiting direct generalization from young to elderly adults.

Beyond local electrophysiological effects around stimulation events, some studies have reported broader changes in oscillatory activity across stimulation periods. These effects may also vary with age: Schneider et al. (2020), for example, found increased SO spectral power in young but not middle-aged adults. However, most previous work has focused on oscillation amplitude, spectral power, spindle activity, or event-related responses time-locked to stimulation. Although temporal relationships between successive SOs have also been examined (Schneider et al., 2020), the effects of CLAS on SO train initiation, persistence, and train-length distributions remain insufficiently characterized.

Therefore, we investigated whether CLAS could help restore rhythmic SO dynamics in elderly adults. To this end, we compared changes from a baseline night to an experimental (EXP) night between a control (SHAM) group without auditory stimulation (N = 11) and a stimulation (STIM) group receiving CLAS (N = 16). Using the temporal classification framework previously introduced by our group (Capurro et al., 2026), we distinguished isolated SOs from trains of consecutive SOs based on inter-SO intervals. SOs were classified as isolated when separated from both neighboring SOs by more than 2 s, whereas successive SOs separated by less than 2 s were grouped into trains. This threshold corresponds to the upper limit of the SO period within the 0.5-1 Hz frequency range. We quantified the proportions of isolated and consecutive SOs and the probability that a newly occurring SO initiated a train. We also examined mean train length and the composition of train-length distributions. Analyses distinguished local effects in online-detected events from overnight effects in offline-detected NREM SOs. Locally, CLAS increased the proportion of consecutive SOs and the probability of train initiation. Additional analyses indicated greater increases in mean train length in both local and global analyses, without statistically significant differential changes in NREM SO density.

## Results

### CLAS enhances local SO train initiation and persistence in online-detected events during detection and stimulation periods

We first investigated the local effects of CLAS on SO temporal organization using the set of online detections generated by the closed-loop system. After restricting events to offline-scored NREM sleep and applying offline waveform reconstruction and quality-control procedures to identify valid SOs and accurately locate their negative peaks (see Methods), temporal organization metrics were computed using the framework introduced in Capurro et al. (2026). SOs were classified as isolated or consecutive according to their temporal proximity to neighboring SOs, and SO trains were identified as sequences of consecutive SOs and characterized by their lengths, defined as the number of SOs within each train. In addition, the first SO of each train was termed a train-start SO. **Figure 1** provides a schematic illustration of the SO classification procedure and the temporal organization metrics derived from it. Changes between the EXP and BASELINE nights were quantified and compared between groups.

**Figure 1.**
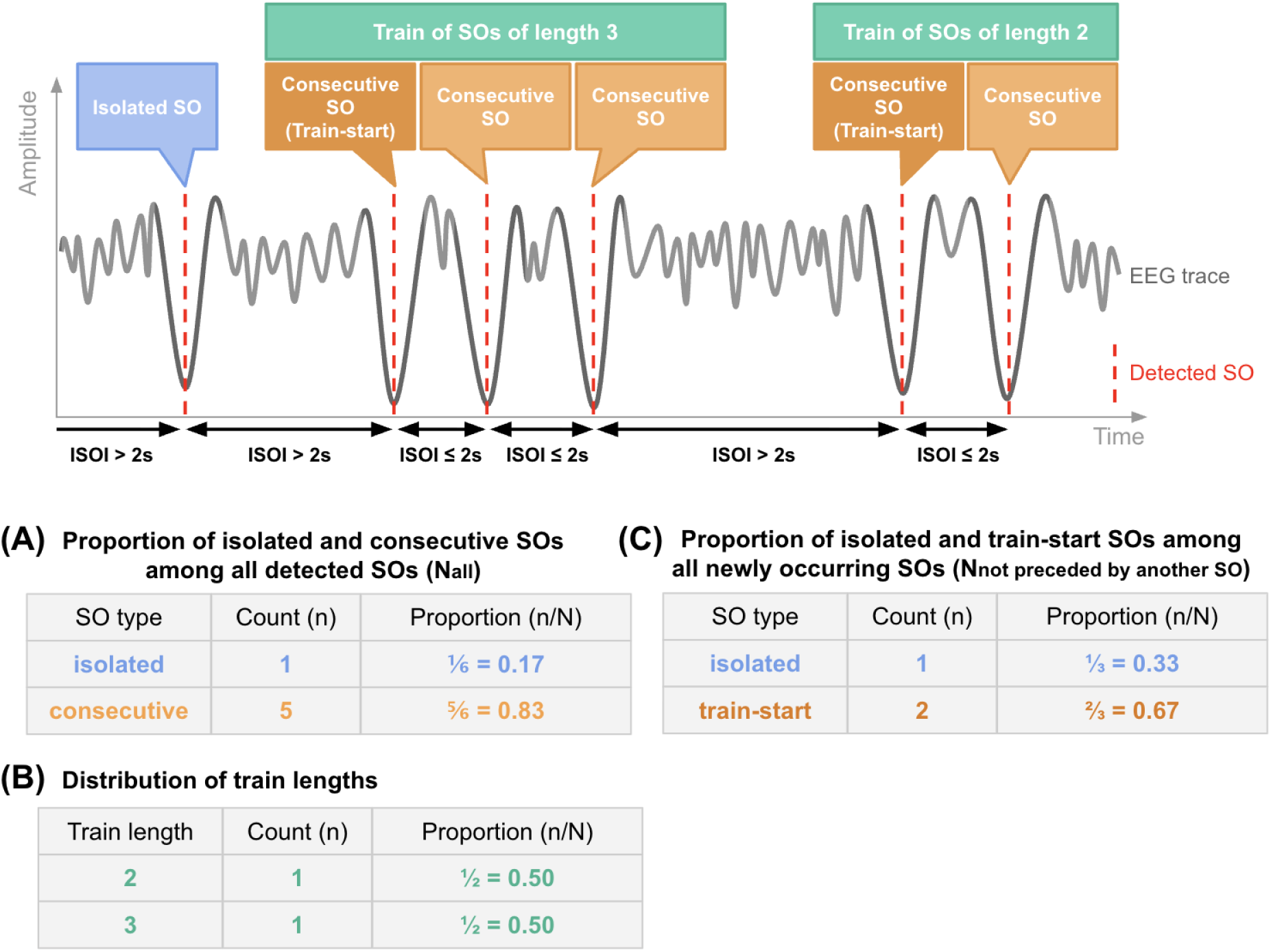
Schematic illustration of the slow oscillation (SO) temporal organization framework. Detected SOs were classified according to the inter-SO interval (ISOI) separating consecutive negative peaks. SOs preceded and followed by intervals greater than 2 s were classified as isolated, whereas SOs occurring within sequences of events separated by ISOIs ≤ 2 s were classified as consecutive. Consecutive SOs formed SO trains, whose length was defined as the number of SOs contained within each train. The first SO of each train was labeled a train-start SO. The lower panels illustrate the three temporal organization metrics derived from this classification: **(A)** the proportion of isolated and consecutive SOs among all detected SOs, **(B)** the distribution of SO train lengths, and **(C)** the proportion of isolated and train-start SOs among all newly occurring SOs (i.e., SOs not preceded by another SO within 2 s). The numerical values shown are illustrative examples.

We observed that CLAS promoted the organization of locally detected SOs into trains, with a greater increase in the proportion of consecutive SOs and a complementary decrease in isolated SOs in STIM compared with SHAM from BASELINE to EXP (group x night interaction: b = 1.209, z = 2.555, p = 0.011; **Fig. 2A**). Among newly occurring SOs, CLAS also increased the probability of initiating a train rather than remaining isolated (group x night interaction: b = 1.093, z = 2.536, p = 0.011; **Fig. 2B**). Both effects remained significant after accounting for SO density (consecutive SOs: b = 0.876, z = 2.077, p = 0.038; train-start SOs: b = 0.775, z = 1.975, p = 0.048).

**Figure 2.**
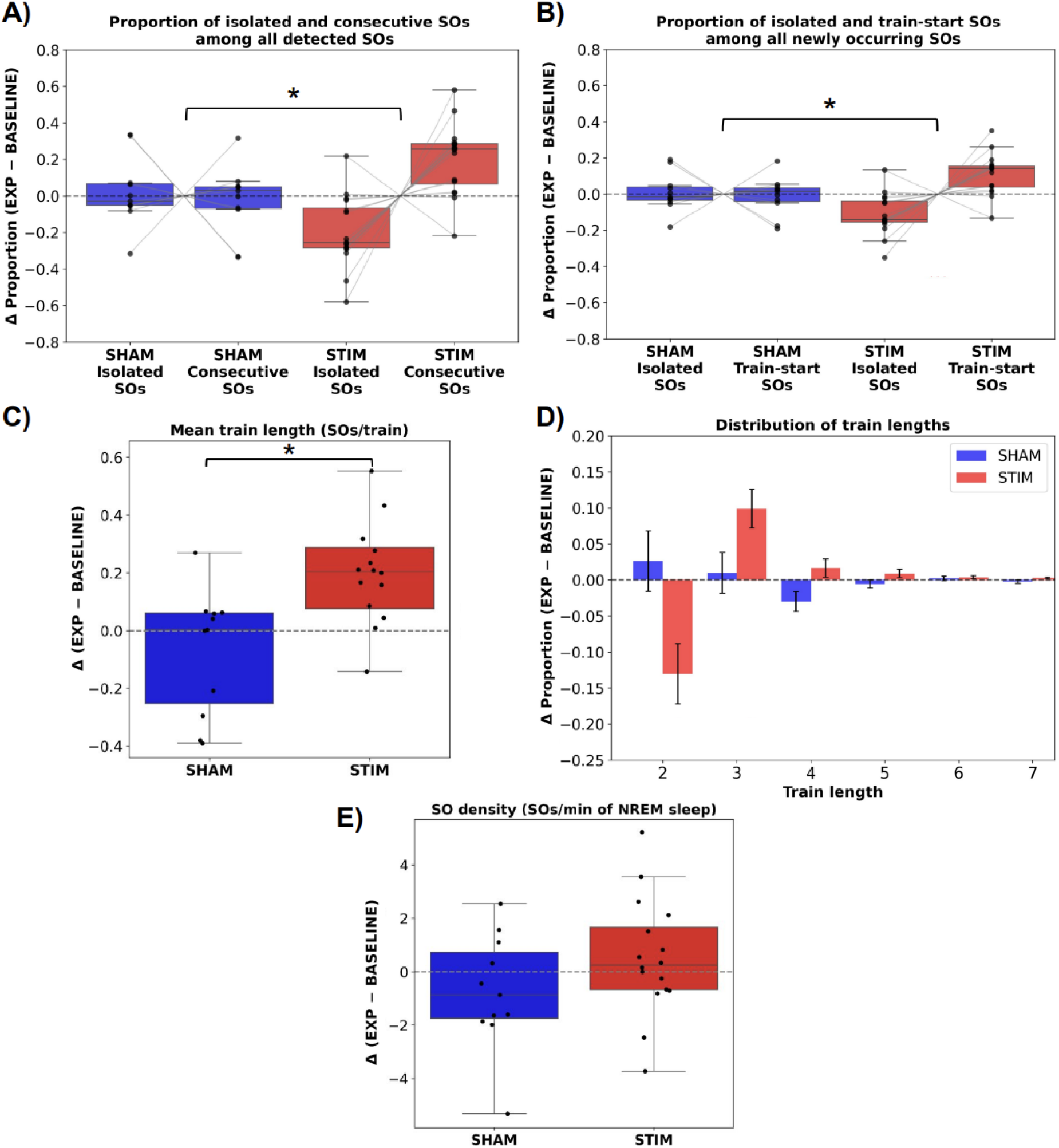
Local effects of CLAS on slow oscillation temporal organization. Within-subject changes between the experimental and baseline nights (EXP - BASELINE) are shown for SHAM (blue, n = 11) and STIM (red, n = 16). **(A)** Proportions of isolated and consecutive SOs among all detected SOs. **(B)** Proportions of isolated and train-start SOs among newly occurring SOs. **(C)** Mean train length, calculated as the proportion-weighted mean number of SOs per train. **(D)** Changes in train-length proportions, shown as group means ± SEM. **(E)** Changes in NREM SO density. Positive values indicate increases from BASELINE to EXP. Individual dots represent subjects. ***p < 0.001; **p < 0.01; *p < 0.05.

We next examined whether this greater propensity to form trains was accompanied by an increase in train length. The mean train length increased more from BASELINE to EXP in STIM than in SHAM (ΔSTIM = 0.184 ± 0.067 vs ΔSHAM = −0.070 ± 0.065 SOs per train; Welch’s t(24.34) = 2.727, p = 0.012; **Fig. 2C**). Consistent with this finding, the distribution of train lengths showed a descriptive shift toward longer trains in STIM relative to SHAM (**Fig. 2D**). However, the joint analysis of the complete train-length composition did not reach statistical significance (ILR-MANOVA: Pillai’s trace = 0.334, F(5,21) = 2.109, p = 0.104). Exploratory component-specific comparisons suggested a relative decrease in two-wave trains and a relative increase in four-wave trains in STIM compared with SHAM, each evaluated against the geometric mean of the remaining train-length proportions. These comparisons reached nominal significance but did not survive Holm correction (two-wave trains: F(1,25) = 7.294, p = 0.012, adjusted p = 0.073; four-wave trains: F(1,25) = 6.593, p = 0.017, adjusted p = 0.083). These changes occurred without a statistically significant between-group difference in the change in NREM SO density (group x night interaction: b = 1.259, t(25) = 1.479, p = 0.152; **Fig. 2E**). Complete statistical results are provided in Supplementary **Table S1**. Together, these findings support a reorganization of local SO activity: a greater proportion of SOs participated in trains, newly occurring SOs were more likely to initiate trains, and trains became longer on average.

### Control analyses support the robustness of local CLAS effects

To determine whether the observed differences in SO temporal organization reflected genuine changes in SO rhythmicity rather than methodological factors, we performed a series of control analyses (see Supplementary Materials).

Control analyses supported the robustness of the local CLAS effects to potential methodological confounds. The temporal distribution of detections and stimulations across the night was comparable across groups and nights, and the increases in consecutive SO proportions, train-initiation probability, and mean train length remained significant without waveform-based rejection. No significant group-by-night interactions were observed in wave morphology or the proportion of faster delta events. These findings suggest that the enhanced SO train initiation and persistence in the STIM group were unlikely to be explained by detection timing, waveform-based rejection, or changes in detected-wave characteristics. Detailed analyses are reported in the Supplementary Materials (**Figs. S1-S5; Tables S3-S4**).

### Increased mean SO train length extends to offline-detected events across the night

We next investigated whether the changes in SO temporal organization observed around stimulation periods were also present at the global overnight level. Because local analyses were restricted to online-detected SOs, train identification could be affected by detector on/off periods, post-stimulation pause intervals, and preprocessing-related event rejection. To assess SO organization independently of these sampling constraints, temporal organization metrics were recomputed using all offline-detected NREM SOs across the full night (see Methods).

Given the local and intermittent nature of stimulation, we expected its effects on SO temporal organization to be attenuated when assessed across the full night. The global analysis did not reveal a significant differential change between STIM and SHAM in the proportion of consecutive SOs (group x night interaction: b = 0.109, z = 0.538, p = 0.590; **Fig. 3A**) or in the probability that a newly occurring SO initiated a train rather than remaining isolated (b = −0.006, z = −0.038, p = 0.970; **Fig. 3B**). Both interactions remained non-significant after accounting for SO density (consecutive SOs: b = 0.222, z = 1.127, p = 0.260; train-start SOs: b = 0.069, z = 0.419, p = 0.676).

**Figure 3.**
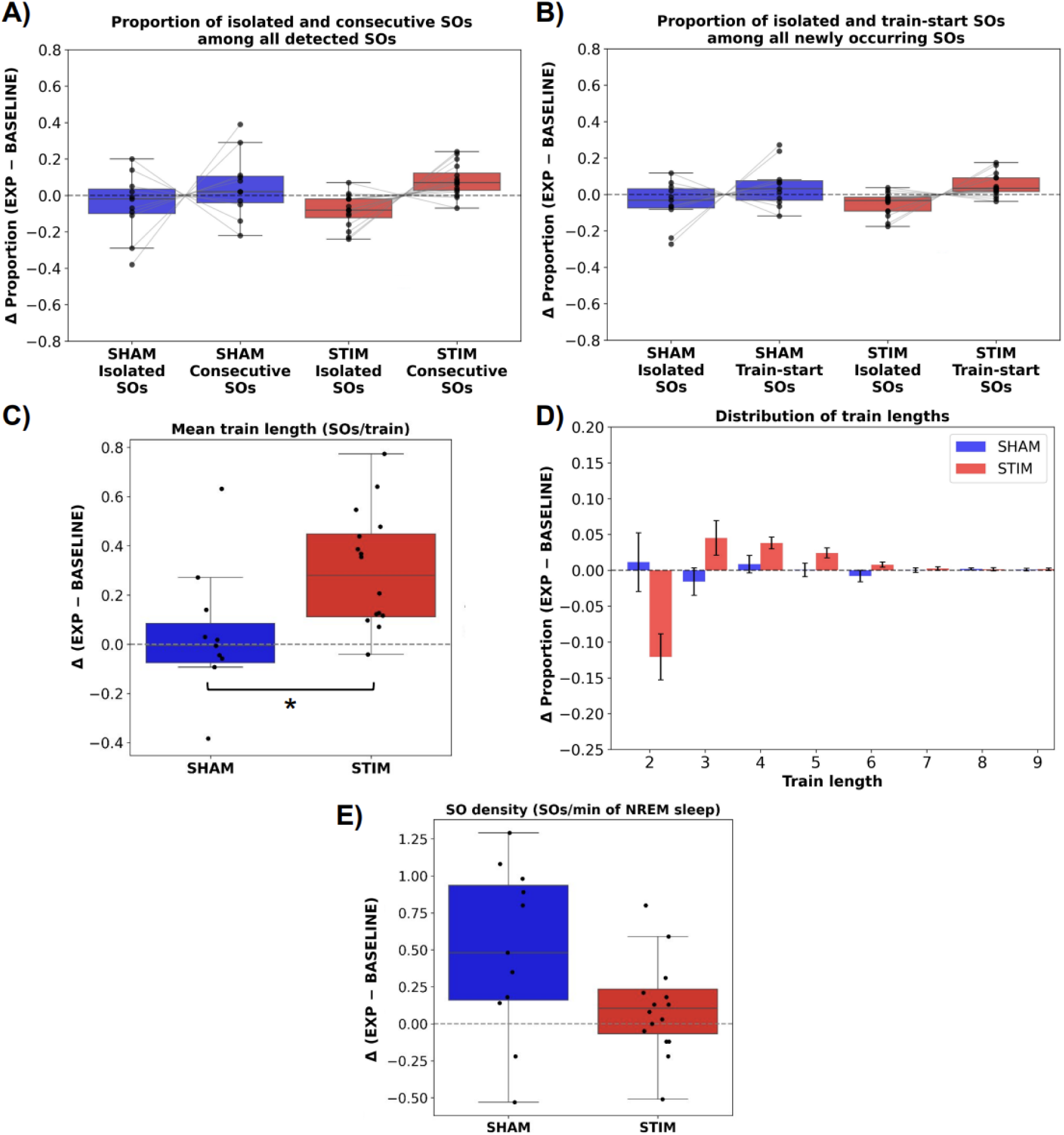
Global effects of CLAS on slow oscillation temporal organization. Within-subject changes between the experimental and baseline nights (EXP - BASELINE) are shown for SHAM (blue, n = 11) and STIM (red, n = 16). **(A)** Proportions of isolated and consecutive SOs among all detected SOs. **(B)** Proportions of isolated and train-start SOs among newly occurring SOs. **(C)** Mean train length, calculated as the proportion-weighted mean number of SOs per train. **(D)** Changes in train-length proportions, shown as group means ± SEM. **(E)** Changes in NREM SO density. Positive values indicate increases from BASELINE to EXP. Individual dots represent subjects. ***p < 0.001; **p < 0.01; *p < 0.05.

Nevertheless, the mean train length increased more from BASELINE to EXP in STIM than in SHAM (ΔSTIM = 0.257 ± 0.079 vs ΔSHAM = −0.010 ± 0.096 SOs per train; Welch’s t(21.46) = 2.138, p = 0.044; **Fig. 3C**). Consistent with this finding, the train-length distribution showed a descriptive shift toward longer trains in STIM relative to SHAM (**Fig. 3D**). However, the joint analysis of the complete train-length composition did not reach statistical significance (ILR-MANOVA: Pillai’s trace = 0.185, F(7,19) = 0.616, p = 0.736). None of the exploratory comparisons of individual train-length categories relative to the remaining categories reached nominal significance or survived Holm correction (all unadjusted p ≥ 0.166; all adjusted p = 1.000). These findings were not accompanied by a statistically significant between-group difference in the change in global NREM SO density (group x night interaction: b = −0.311, t(25) = −1.560, p = 0.131; **Fig. 3E**). Complete statistical results are provided in Supplementary **Table S2**. Together, these findings suggest that the increase in average SO train length extended to overnight recordings, whereas corresponding changes in the proportion of SOs participating in trains or initiating trains were not statistically detectable at the global level.

### Global sleep structure and spectral composition are comparable between groups

To obtain a comprehensive overview of sleep patterns in both groups, we performed conventional analyses of sleep architecture and power spectral density. To date, no published study has shown that a single night of CLAS is sufficient to induce changes in sleep architecture or spectral power across the entire night (Ngo et al., 2013, 2015; Golrou et al., 2018; Schneider et al., 2020; Wunderlin et al., 2023; Zeller et al., 2024). Therefore, although no major stimulation-induced changes in overall sleep structure were expected, we performed standard analyses to characterize sleep patterns in both groups.

Most sleep architecture measures showed no significant differential changes between the SHAM and STIM groups from BASELINE to EXP (**Table S6**). These included total sleep time (TST), time spent awake after sleep onset and in N1, N2, N3, and NREM sleep, overall sleep-stage composition, the relative proportions of N2 and N3 within NREM, and measures of sleep fragmentation. REM sleep duration was the only measure showing a significant group × night interaction (F(1, 25) = 5.81, p = 0.024), with a significant increase from BASELINE to EXP in SHAM but no significant change in STIM. Sleep latency showed a significant main effect of night (F(1, 25) = 8.17, p = 0.008), decreasing during EXP in both groups, consistent with habituation to the laboratory environment. Relative power spectral density analyses across sleep stages also revealed no significant effects of stimulation (**Table S7**), with no significant group differences or group x night interactions detected for any frequency band.

Thus, consistent with previous CLAS studies, a single night of stimulation did not substantially alter sleep architecture or the overall spectral composition of sleep, and both groups exhibited broadly comparable sleep profiles, indicating that the differences in SO temporal organization were not secondary to changes in global sleep structure.

## Discussion

Locally, stimulation increased the proportion of consecutive SOs and the probability that a newly occurring SO initiated a train, with both effects remaining significant after accounting for SO density. Greater increases in mean train length in STIM than in SHAM were also observed both locally and across overnight NREM recordings, although corresponding changes in consecutive and train-start SO proportions were not detected globally. These findings occurred without statistically significant between-group differences in changes in NREM SO density or relative spectral power, while most sleep architecture measures also showed no significant differential changes. Together, these findings support a reorganization of the temporal structure of SO activity, characterized by enhanced local train formation and increased average train length.

The present findings extend previous work on acoustic stimulation during sleep. Most CLAS studies have focused on changes in SO amplitude, slow-wave activity, spindle activity, SO-spindle coupling, or stimulation-locked responses (Ngo et al., 2013, 2015; Ong et al., 2016; Papalambros et al., 2017; Wunderlin et al., 2021). In contrast, relatively little attention has been paid to the temporal organization of SO activity itself. Recently, our group reported that aging is associated not only with reductions in SO amplitude and density but also with a marked disruption of SO temporal organization, characterized by an increased prevalence of isolated SOs and shorter SO trains (Capurro et al., 2026). The present results indicate that CLAS partially reverses these age-related alterations by promoting the generation and persistence of rhythmic SO sequences. These findings suggest that acoustic stimulation may restore aspects of SO temporal organization that are disrupted with aging, extending its effects beyond previously described changes in SO amplitude and related oscillatory measures.

The absence of significant changes in overall SO density is particularly interesting. Many studies evaluating CLAS efficacy have emphasized increases in slow-wave activity or the number of detected oscillations as primary outcomes. However, the present findings suggest that stimulation can modify the temporal arrangement of SOs even when the total number of events during NREM sleep remains unchanged. This dissociation implies that the physiological relevance of slow-wave activity may depend not only on how many SOs occur, but also on how they are organized across time. From this perspective, temporal organization metrics may capture aspects of sleep physiology that remain invisible to traditional density- or amplitude-based analyses.

Although the functional significance of SO trains remains largely unexplored, several lines of evidence suggest that sustained rhythmic activity may be particularly relevant for sleep-dependent brain functions. Recent evidence indicates that rhythmic neural activity during sleep contributes to glymphatic transport and metabolic waste clearance (Fultz et al., 2019; Jiang-Xie et al., 2024). In this context, repeated oscillatory cycles may provide a more effective driving mechanism for cerebrospinal fluid movement than isolated events. Although the present study did not directly assess memory performance or glymphatic function, our findings raise the possibility that restoring rhythmic SO persistence may represent an additional mechanism through which CLAS exerts beneficial physiological effects.

An important aspect of the present study is the convergence between local and global analyses for increased mean train length. The observation of comparable effects using two independent detection approaches strengthens the interpretation that CLAS genuinely enhances SO rhythmicity rather than producing an artifact of the online detection procedure.

Moreover, several control analyses argue against alternative methodological explanations for the observed effects. First, the temporal distribution of detections across the night was highly comparable between nights and groups. If anything, the small shifts observed across nights would be expected to favor greater SO organization in the SHAM group and reduced organization in the STIM group, opposite to the observed pattern. Second, the results remained unchanged when waveform-based rejection procedures were omitted, indicating that they do not arise from preprocessing decisions. Third, neither local nor global analyses revealed stimulation-related changes in conventional SO morphological characteristics, suggesting that the findings cannot be explained by differences in wave detectability or waveform shape. Finally, the slight decrease in frequency observed across nights occurred similarly in both groups and would theoretically be expected to reduce, rather than increase, the probability of consecutive events because longer periods increase the intervals between successive SOs. Collectively, these observations indicate that potential methodological confounds would tend to attenuate rather than inflate the observed stimulation effects. Another methodological aspect that deserves consideration is the one-second pause period imposed after each stimulation (see Methods). Because no additional detections could be generated during this interval, the pause artificially reduces the probability of detecting closely spaced SOs and can fragment trains that would otherwise continue uninterrupted. Consequently, the pause period introduces a bias against the detection of increased rhythmicity. The fact that stimulation-related increases in train initiation and persistence were observed despite this constraint suggests that the true physiological effects may be larger than those estimated by the local analyses.

Another important limitation concerns the temporal precision of the stimulation platform. Although stimulation was intended to target the SO up-state, the OpenBCI-based system introduced both a delay and timing jitter due to its block-based data acquisition, software processing, operating-system scheduling, and audio-output pipeline, preventing accurate determination of the true auditory stimulation onset from the stored stimulation markers (Ramele et al., 2025). Previous studies have reported temporal offsets and variability in OpenBCI-based EEG recordings, highlighting the influence of buffering, data-processing configurations, and timestamp correction procedures on temporal accuracy (Frey, 2016; Pattisapu and Ray, 2023; Qiu et al., 2019; Knierim et al., 2023; Cardona-Álvarez et al., 2023). Consequently, in our set up, stimulation may have occurred at different phases of the ongoing SO across events. Some stimuli likely occurred before the intended target, others near the optimal up-state region, and others later on. Therefore, the present study cannot determine which stimulation phase was most effective for promoting SO train initiation and persistence. Importantly, this temporal variability would be expected to reduce overall stimulation efficacy by diminishing phase-specific effects. Despite this limitation, robust increases in SO train initiation and persistence were still observed. This suggests that the reported effects are sufficiently strong to emerge even under conditions of substantial phase uncertainty and raises the possibility that greater effects could be achieved using systems with improved temporal precision. Beyond this study-specific limitation, these findings highlight a critical methodological consideration for future CLAS research, particularly in elderly adults. Previous work has shown that the effective phase window for auditory stimulation is narrower in elderly than in younger adults, with optimal effects occurring near the SO positive peak (Navarrete et al., 2020). In this context, both the systematic latency and, more importantly, the temporal jitter of stimulation devices become critical determinants of phase-targeting accuracy. While a stable delay can be characterized and compensated for, trial-to-trial variability in auditory onset introduces phase uncertainty that cannot be corrected by a fixed temporal adjustment. Therefore, consistent with recommendations for methodological standardization in CLAS research (Esfahani et al., 2023), we argue that stimulation latency and jitter should be systematically measured and explicitly reported in future studies. This is particularly important when translating CLAS protocols to elderly populations, in whom the narrower effective stimulation window may make treatment efficacy especially sensitive to temporal variability.

Beyond its implications for stimulation phase targeting, classical event-related analyses time-locked to stimulation could not be reliably performed. This limitation is particularly relevant because stimulation-induced ERP responses during CLAS are typically small in amplitude, often on the order of only 5-10 μV (Krugliakova et al., 2020). Under these conditions, temporal variability in stimulation timing can substantially attenuate averaged responses and markedly reduce sensitivity to stimulation-induced effects. Thus, the absence of ERP analyses should not be interpreted as evidence against stimulation efficacy but rather as a consequence of insufficient temporal precision for detecting subtle phase-locked responses. Importantly, the temporal organization metrics employed here are inherently less sensitive to stimulation latency and jitter because they quantify changes in ongoing oscillatory dynamics over longer timescales rather than relying on millisecond-level temporal alignment. In this sense, the present findings highlight the potential value of temporal organization measures as robust markers of stimulation efficacy in systems where precise timing information is unavailable.

Additional limitations should also be considered. The sample size was relatively modest, and subjects were retrospectively selected from a larger dataset according to stimulation and signal-quality criteria. Moreover, although CLAS altered the temporal dynamics of SO activity, the study did not include cognitive, behavioral, or physiological outcome measures. Consequently, the functional significance of the observed electrophysiological changes remains to be established.

Future studies should involve stimulation platforms with lower latency and reduced temporal jitter to investigate the phase dependence of SO train formation and persistence. Determining whether specific SO phases preferentially promote train initiation and persistence represents an important next step. Previous CLAS studies have identified the SO up-state as an effective target for enhancing stimulation-evoked responses and slow-wave amplitude (Ngo et al., 2013; Ong et al., 2018; Wunderlin et al., 2021), but it remains unknown whether the same phase optimally promotes the emergence of extended SO trains. Indeed, the stimulation phase that maximizes individual SO enhancement or ERP amplitude may not necessarily coincide with the phase that most effectively sustains rhythmic slow oscillations. Addressing this question could help disentangle the mechanisms underlying SO train persistence and contribute to the optimization of CLAS protocols in aging populations. Improved temporal precision would also enable more reliable evaluation of stimulation-locked responses, SO-spindle interactions, and their relationship with changes in SO temporal organization, providing a more comprehensive understanding of the mechanisms through which CLAS influences sleep physiology. Beyond mechanistic studies, future work should determine whether stimulation-induced increases in SO train persistence translate into improvements in glymphatic clearance, learning, memory consolidation, or executive function, or into favorable changes in biomarkers of brain health. Finally, longitudinal interventions will be necessary to establish whether repeated enhancement of SO rhythmicity can produce long-lasting physiological or cognitive benefits in elderly adults.

In conclusion, CLAS promoted local SO train formation in elderly adults, with a greater increase in mean train length also observed across overnight NREM recordings. These effects occurred without statistically significant between-group differences in changes in NREM SO density, and the local increases in consecutive and train-start SO proportions remained significant after accounting for SO density. Most sleep architecture measures and relative spectral power likewise showed no significant differential changes. Together, these findings suggest that CLAS can modify a dimension of SO temporal organization previously shown to be disrupted during aging. More broadly, these findings suggest that understanding how SOs are organized across time may be as important as quantifying how many occur, opening new avenues for the study and modulation of sleep physiology in aging.

## Methods

### Subjects

Subjects were retrospectively selected from a larger dataset of overnight sleep recordings from the Laboratorio de Sueño y Memoria at the Instituto Tecnológico de Buenos Aires (ITBA) acquired as part of a study investigating the effects of CLAS in elderly adults. Elderly adults aged 60-80 years were recruited through the lab’s social media platforms and were screened to exclude severe medical, neurological, or psychiatric conditions.

All control subjects who did not receive any acoustic stimulation were included in the present study to form the control (SHAM) group, while stimulated subjects were selected based on stimulation parameters used during the experimental sessions to form the stimulated (STIM) group. Because the OpenBCI-based stimulation platform introduced latency and timing jitter between the stimulation trigger and the actual acoustic onset (Ramele et al., 2025), consistent with timing limitations previously reported in OpenBCI-based EEG recordings (Frey, 2016; Pattisapu and Ray, 2023; Qiu et al., 2019; Knierim et al., 2023; Cardona-Álvarez et al., 2023) and auditory stimulation systems (Corneyllie et al., 2021), only subjects with a programmed stimulation delay of ≤60 ms after the positive-going zero-crossing were included in the STIM group (see EEG Acquisition System section).

This criterion was adopted to minimize deviations from the intended stimulation target within the SO up-state (Ngo et al., 2013), taking into account the additional latency and temporal variability introduced by the stimulation system. From this subset, two subjects were excluded from further analyses due to poor signal quality, incomplete recordings, or technical issues affecting data acquisition. Thus, a total of 27 elderly subjects were included in the present study: 16 subjects (66.44 ± 1.32 years; 12 women, 4 men) were assigned to the STIM group and received acoustic stimulation during the experimental night, whereas 11 subjects (68.73 ± 1.59 years; 9 women, 2 men) were assigned to the SHAM group and underwent the same experimental protocol without auditory stimulation.

All subjects provided written informed consent prior to participation. The study protocol was reviewed and approved by the Comité de Ética Humana, Facultad de Ciencias Médicas, Universidad de Buenos Aires. Participants received financial compensation for their participation, and transportation to and from the study site was provided.

### Experimental Design

Subjects underwent two overnight sleep recordings separated by one week: an adaptation night and an experimental night. Although originally intended as an adaptation night, it was subsequently used as the baseline (BASELINE) for the present analyses. Therefore, no separate adaptation night was available before baseline assessment. Both recording sessions were conducted in the sleep laboratory under identical environmental conditions between approximately 21:00 and 07:30 h.

During the baseline night, sleep stages were monitored and slow oscillations were detected online while subjects slept under conditions identical to those of the EXP night but without acoustic stimulation. This recording allowed the characterization of baseline sleep physiology and the analysis of changes relative to this baseline, thereby accounting for inter-subject variability. In addition, an individual auditory stimulation threshold was determined for each subject. After the first sleep spindle was detected online and stable N2 sleep had been maintained for approximately 10 minutes, some auditory stimuli were presented to identify the highest stimulation intensity that did not induce awakenings or visible arousals.

Subjects returned to the laboratory seven days later for the EXP night. The recording setup and online detection procedures were identical to those used during BASELINE. Subjects were assigned to either the SHAM or STIM group. In the STIM group, auditory stimuli were delivered using a CLAS protocol targeting the up-state of ongoing SOs according to the stimulation parameters determined during the baseline night. In the SHAM group, the online detection algorithm operated identically to the BASELINE, with no auditory stimuli being presented.

Analyses focused on the change between the experimental and baseline nights (EXP - BASELINE) for each subject. Differences in these night-to-night changes were then compared between the SHAM and STIM groups to isolate stimulation-related effects from spontaneous variability occurring across recording sessions.

### EEG Acquisition System

Polysomnographic recordings including electroencephalographic (EEG), electrooculographic (EOG), and chin electromyographic (EMG) signals were acquired using a custom real-time sleep monitoring and stimulation platform based on Cyton and Daisy biosignal amplifiers (OpenBCI, USA). The acquisition system consisted of an Ubuntu 22.04 workstation equipped with a low-latency preempt_rt kernel and the jackd2 audio server to improve real-time performance during auditory stimulation. Custom software was developed in Python 3.6.10 using the OpenBCI Python driver for signal acquisition and the alsaaudio package for audio generation.

Signals were sampled at 250 Hz and streamed in real time to the stimulation platform. EEG activity was recorded from F3, F4, C3, C4, P3, and P4 according to the international 10-20 system and referenced to a single mastoid. Two bipolar EOG channels and two bipolar chin EMG channels were also recorded. During online monitoring and stimulation, EEG signals were displayed and processed using a 0.16-30 Hz band-pass filter, EOG signals using a 0.2-35 Hz band-pass filter, and EMG signals using a 10-90 Hz band-pass filter. However, the recorded data were stored in their raw form, with only a 0.16 Hz high-pass filter applied after acquisition to remove slow drifts and keep the signals centered around zero.

Auditory stimulation was delivered through headphones using brief bursts of pink noise presented at the individually adjusted intensity level determined during the BASELINE night. The OpenBCI-based stimulation platform processed incoming EEG data in blocks of 16 samples at a sampling rate of 250 Hz, corresponding to a block duration of 64 ms. If processing awaited the arrival of a complete block, the buffering stage could introduce a sample-dependent waiting time ranging from 0 to 60 ms, with a theoretical mean of 30 ms and a standard deviation of approximately 18.5 ms, assuming that events occurred with equal probability at each sample position within the block. These values represent an idealized estimate of the buffering contribution rather than measurements of the actual system latency or jitter. In practice, additional temporal delays and variability may arise from wireless data transmission, software execution, operating-system scheduling, and audio output.

Previous studies have reported temporal offsets and variability in OpenBCI-based EEG recordings, highlighting the influence of buffering and data-processing configurations, although the magnitude of these offsets depends on the acquisition hardware, software configuration, and timing-correction procedures. Frey (2016) reported an approximately 88-ms offset between event-related potentials recorded with OpenBCI and a reference amplifier, whereas Qiu et al. (2019) measured a mean difference of 47.23 ± 3.56 ms between software-defined stimulus markers and OpenBCI timestamps. Pattisapu and Ray (2023) also compared event-related responses recorded with OpenBCI and a research-grade EEG amplifier. In a distributed OpenBCI framework, Cardona-Álvarez et al. (2023) reported an acquisition-system latency of approximately 56 ms under their specific configuration. Furthermore, Knierim et al. (2023) demonstrated that the timing offset and jitter of OpenBCI Cyton+Daisy recordings can vary substantially with buffer configuration and timestamp-correction methods. In addition, the delay and jitter of auditory stimulation systems can vary substantially depending on their hardware and software configurations (Corneyllie et al., 2021). These findings highlight that timing performance is implementation-dependent and that latency estimates obtained using different measurement procedures are not directly interchangeable.

In our setup, the acquisition buffering, together with software, operating-system, and audio-output delays, resulted in an empirically measured end-to-end stimulation delay of approximately 80 ms (Ramele et al., 2025), while buffering could contribute a sample-dependent variability of up to 60 ms in the worst-case scenario. Consequently, stimulation did not occur immediately when it was triggered. Thus, the stored stimulation markers represent trigger times rather than the exact onset of auditory stimulation. Instead, the true stimulation onset occurred somewhere within a temporal uncertainty window centered around the delay. This limitation is particularly relevant in CLAS paradigms because the optimal stimulation window is restricted to a narrow portion of the SO up-state (Navarrete et al., 2020). Large delays may cause stimulation to occur after the intended target phase, thereby reducing stimulation efficacy. For this reason, only subjects whose stimulation delay parameter was small were included in the present study.

### Online Sleep Staging, Online SO detection and CLAS Protocol

Real-time sleep staging was continuously performed by experimenters through visual inspection of the polysomnographic recordings throughout the night to identify periods of NREM sleep suitable for SO detection and auditory stimulation. SO detection and, in the STIM group, auditory stimulation were manually enabled after 10 min of stable N2 sleep and subsequently restricted to N2 and N3 sleep under continuous experimenter supervision. Because detection and stimulation opportunities were restricted to online-identified NREM sleep periods, we evaluated the temporal distribution of detections and stimulations across the night to ensure that events occurred during comparable portions of the sleep period across nights and experimental groups. This control analysis was particularly relevant because slow-wave activity varies substantially throughout the night (Massimini et al., 2004). These analyses are reported in the Supplementary Materials.

The SO detection algorithm continuously monitored a single EEG channel (C3 or C4) selected according to signal quality. The use of central electrodes for online slow-wave detection is supported by previous CLAS studies demonstrating effective phase-targeted auditory stimulation using C4 (Krugliakova et al., 2020). The target events were SOs in the 0.5-1 Hz range. However, given the real-time constraints of the detection system, the online signal was low-pass filtered at 4 Hz to isolate slow oscillatory activity while preserving the slow-wave morphology relevant to detection. SO detection was implemented using a causal finite-state machine operating on the filtered signal. Detection was based on the occurrence of a negative half-wave followed by a positive-going zero-crossing. Specifically, the algorithm continuously tracked the maximum signal value within a 1-s window and identified candidate events when the signal decreased by at least 60 μV relative to this maximum and crossed a negative threshold of −40 μV.

Once these criteria were fulfilled, a detection marker was generated and stored in the recording. The subsequent positive-going zero-crossing then triggered the stimulation sequence. After a subject-specific predefined stimulation delay was applied in order to target the upcoming positive peak of the SO up-state, auditory stimulation was triggered and a stimulation marker was also stored in the recording. A pause period of 1 s was imposed after each stimulation event, during which no additional SO detections were considered.

Importantly, due to the latency and timing uncertainty of the stimulation platform described above, stimulation markers represent the stimulation trigger rather than the exact onset of auditory stimulation.

### Offline Signal Preprocessing and Sleep Staging

EEG recordings were processed offline using Python (version 3.10.19), primarily with the MNE package (version 1.7.1) (Gramfort et al., 2013).

All offline analyses were performed on a single central EEG channel per recording. Central derivations were chosen as the probability of detecting a slow oscillation is the highest at central and frontal electrodes (Massimini et al., 2004). The best-quality central channel (C3 or C4) was selected for each night of each subject based on visual inspection.

Sleep stages were automatically scored in 30-second epochs using the Yet Another Spindle Algorithm (YASA) sleep staging function (version 0.6.5) (Vallat & Walker, 2021), according to the American Academy of Sleep Medicine (AASM) criteria (Iber et al., 2007). Sleep stages were classified as Wake (W), N1, N2, N3, and REM. Following the recommendations provided in the function’s documentation, sleep staging was performed using the selected EEG channel together with EOG and EMG signals without applying additional filtering prior to classification. Subject age and sex were provided to the algorithm through the optional metadata parameters.

The overall performance of the YASA algorithm has been reported to achieve an accuracy of ∼0.85 compared to expert consensus scoring, matching human inter-rater agreement, with F1-score of ∼0.86 for Wake, N2 and REM sleep, ∼0.84 for N3, and ∼0.43 for N1 (Vallat & Walker, 2021). N1 is the least reliable sleep stage both for automated algorithms and human scorers, reflecting its inherently low inter-rater agreement (<45%, (Malhotra et al.,2013)).

To assess the reliability of automated staging in the present dataset, YASA classifications were compared against expert manual scorings available for a subset of recordings. Consistent with previous validation studies, high agreement was observed for Wake, N2, N3, and REM sleep, whereas N1 showed substantially lower performance. Overall accuracy was 78.1% for the full five-stage classification and 88.6% after grouping stages into Wake/Light Sleep, NREM Sleep, and REM Sleep. Detailed confusion matrices and classification metrics are provided in the Supplementary Materials.

After sleep staging, the selected EEG channel was band-pass filtered between 0.3 and 35 Hz and notch filtered at 50 Hz using zero-phase finite impulse response (FIR) filters prior to further analyses.

Because the wireless acquisition system occasionally lost data packets during transmission, the acquisition drivers interpolated the missing samples. These periods were automatically identified by interpolation markers generated by the acquisition software. Segments containing interpolated samples were treated as artifact-contaminated and excluded from signal-based analyses, and any SO detections occurring within these segments were discarded. In addition, sleep-scoring epochs containing more than 10 s of interpolated samples were considered invalid for staging and were assigned as artifact (ART).

### Local Analysis of Stimulation Effects

Local analyses were performed using the set of slow-wave detections generated online by the closed-loop stimulation system. Because detections were generated in real time from an online-filtered EEG signal and detection markers were placed as soon as the predefined detection criteria were satisfied, additional offline preprocessing was required before analyzing the detected events. This procedure aimed to remove erroneous detections, relocate detection markers to the actual negative peak of the underlying waveform, and reconstruct the complete waveform of each detected event.

First, a small number of detections from the experimental night of the STIM group were excluded because they were not followed by an auditory stimulus. During overnight recordings, both detection and stimulation could be manually enabled or disabled by the supervising technicians. Consequently, a few detections were generated after stimulation had been paused, mainly toward the end of stimulation periods, and were therefore not considered for stimulation-related analyses. Second, detections were restricted to NREM sleep. Given the high agreement observed between automatic and manual sleep staging for NREM sleep (see Supplementary Materials, **Fig. S9** and **Table S9**), all detections occurring outside offline-scored N2 or N3 sleep were discarded.

The remaining detections were then inspected offline to reconstruct the underlying waveform morphology (**Fig. 4**). Because the online detector operated causally and relied on simplified criteria optimized for rapid stimulation delivery, detection markers did not necessarily coincide with the true negative peak of the slow wave. Furthermore, some detections did not correspond to genuine slow waves when the complete EEG waveform became available for inspection.

**Figure 4.**
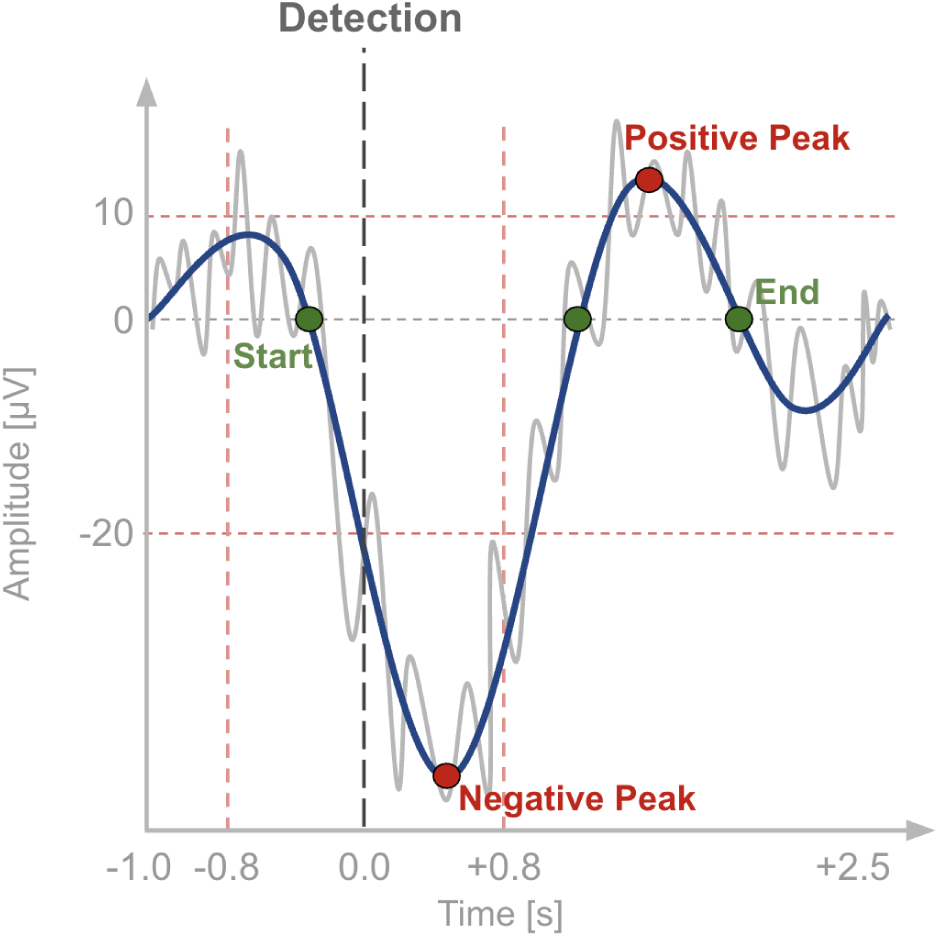
Schematic representation of the procedure used to reconstruct slow-wave morphology and identify characteristic waveform landmarks. The online detection marker (black dashed line) indicates the time at which the real-time detection criteria were satisfied. Offline processing identified the corresponding negative and positive peaks (red), and the zero crossings (green), thereby defining a complete slow-wave cycle. Gray trace: preprocessed EEG signal. Blue trace: 0.5-4 Hz filtered signal.

For each detection, a segment extending from 1 s before to 2.5 s after the original detection marker was extracted from the selected EEG channel. The signal was filtered using a zero-phase fourth-order high-pass filter at 0.5 Hz followed by a zero-phase fourth-order low-pass filter at 4 Hz. To compensate for amplitude attenuation introduced by filtering, the filtered signal was rescaled to match the standard deviation of the high-pass filtered signal. Candidate negative peaks were identified as prominent negative deflections detected within a temporal window centered on the original detection marker. Peaks were required to exhibit a minimum prominence of 30 μV, a minimum width of 10 samples, and an amplitude lower than −20 μV. Among all valid candidates, the negative peak closest to the original detection marker was selected. Detections for which no valid negative peak was identified within ±0.8 s of the online marker were rejected. For each candidate event, the preceding negative-going zero crossing and the subsequent positive-going zero crossing were identified. A positive peak was then searched after the positive-going zero crossing and was required to exceed +10 μV. Finally, the subsequent descending zero crossing was identified, defining the end of the positive half-wave. After the cycle boundaries had been determined, the negative and positive peaks were redefined as the minimum and maximum amplitudes, respectively, within the reconstructed waveform. Specifically, the negative peak was defined as the minimum value between the initial and final zero crossings, whereas the positive peak was defined as the maximum value between the positive-going and descending zero crossings. Together, these landmarks delineated a complete slow-wave cycle (**Fig. 4**).

Detections for which any of these characteristic waveform landmarks could not be reliably identified were considered erroneous online detections and were excluded from local analyses. The proportion of detections excluded at each preprocessing step is reported in the Supplementary Materials (**Fig. S2**). Because these preprocessing steps reduced the number of retained detections, all analyses of detection outcomes were expressed as proportions relative to the total number of valid detections, allowing direct comparisons across subjects and groups despite differences in the number of retained events. To evaluate whether the main findings depended on the waveform-based cleaning procedure, all primary analyses were repeated using the complete set of NREM detections without morphological rejection. These control analyses are also reported in the Supplementary Materials (**Fig. S3**).

### Temporal Organization Analysis

Temporal organization of SOs was quantified using the methods described in Capurro et al. (2026). SOs were classified as isolated or consecutive according to the inter-SO intervals (ISOIs) measured between markers located at the negative peaks of each SO. SO trains were defined as sequences of consecutive SOs, and train length was defined as the number of SOs contained within each sequence. The first SO of each train was termed a train-start SO.

For each night and group, we quantified the proportions of isolated and consecutive SOs relative to the total number of detected SOs and the distribution of train lengths. To assess whether SO trains became longer on average, mean train length was calculated for each subject and night by weighting each train length by its corresponding proportion and summing across all categories. All available train-length categories were included: 2-7 SOs for the local morphology-cleaned and morphology-unfiltered datasets, and 2-9 SOs for the global dataset. In addition, we focused on SOs that were not preceded by another consecutive SO (i.e., events that could either remain isolated or initiate a train). Within this subset, we computed the proportions of isolated SOs and train-start SOs. This metric reflects the probability that a newly occurring SO either remains an isolated event or recruits subsequent SOs, giving rise to a train.

Because the total number of detected SOs could vary across nights, subjects, and experimental groups (as a consequence of physiological differences in slow-wave activity as well as differences in detection opportunities and preprocessing-related event rejection) these measures were expressed as proportions. This normalization allowed comparisons of temporal organization that were independent of differences in the absolute number of detections. Differences between BASELINE and EXP nights were then compared between the SHAM and STIM groups.

### SO features

To determine whether stimulation altered the morphology of individual SOs, a set of conventional morphological features was extracted from each event. Feature extraction was based on the waveform morphological characteristics identified during the offline verification procedure described above, namely the negative peak, positive peak, initial zero-crossing, mid zero-crossing, and final zero-crossing (**Fig. 5**).

**Figure 5.**
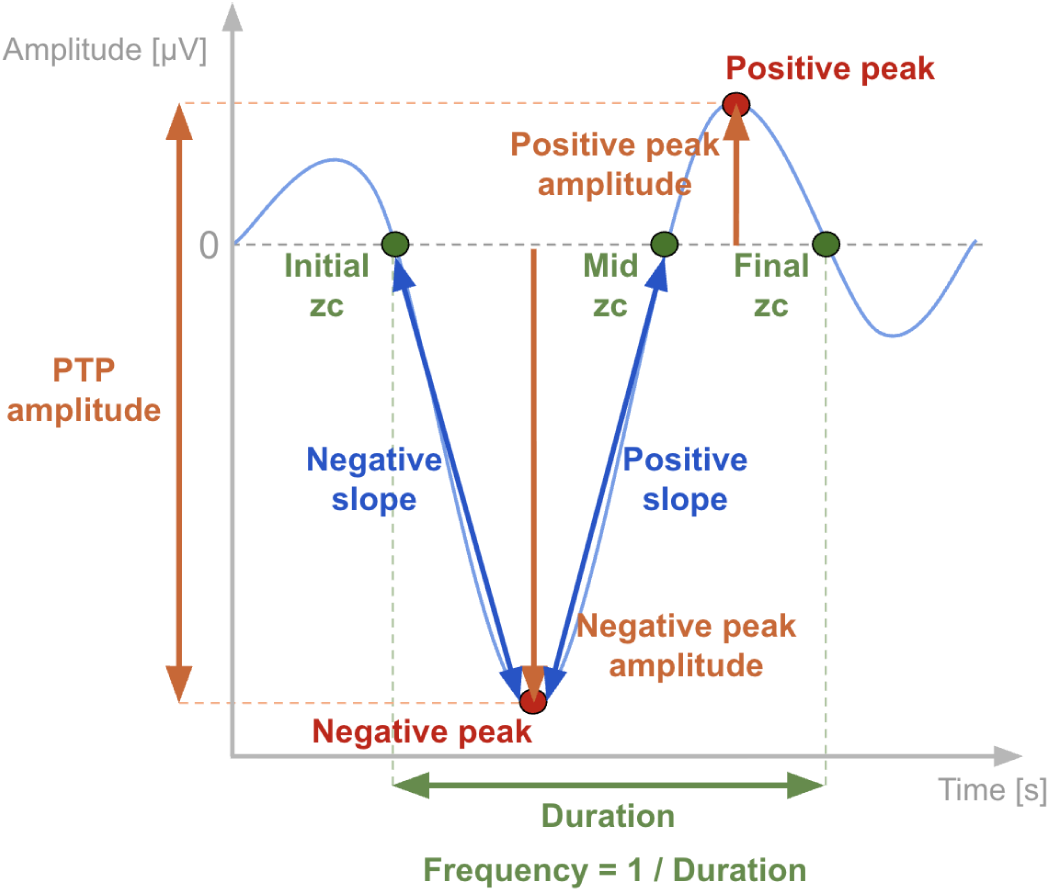
Schematic representation of the procedure used to compute waveform features from characteristic waveform landmarks. The initial, mid, and final zero crossings (zc) define the slow-wave cycle and are used together with the negative and positive peaks to compute negative and positive peak amplitudes, peak-to-peak (PTP) amplitude, event duration, frequency, and positive and negative slopes. Duration was defined as the interval between the initial and final zero crossings, frequency as its inverse, negative slope from the initial zero crossing to the negative peak, and positive slope from the negative peak to the mid zero crossing. Feature extraction was performed on the 0.5-4 Hz filtered waveform.

For each slow wave, the negative peak amplitude and positive peak amplitude were obtained from the corresponding values of the waveform at the location of the peaks in the 0.5-4Hz band-pass filtered and scaled signal. Peak-to-peak amplitude was calculated as the difference between the positive and negative peak amplitudes. Total event duration was defined as the interval between the initial and final zero crossings, and event frequency was estimated as the inverse of this duration. The negative slope was defined as the negative peak amplitude divided by the time interval between the initial zero crossing and the negative peak. The positive slope was defined as the absolute amplitude of the negative peak divided by the time interval between the negative peak and the mid zero crossing. For each recording, feature values were averaged across all validated slow waves, yielding a single value per subject and night. These night-level averages were subsequently used for statistical analyses.

Because online detection relied on a broad low-pass filter and was optimized for real-time stimulation rather than strict waveform classification, validated events were not necessarily restricted to slow oscillations (≤1 Hz) and could also include faster delta waves. Therefore, the frequency of each reconstructed event was examined to characterize the proportion of detections corresponding to slow oscillations and delta activity. These analyses are reported in the Supplementary Materials (**Fig. S5**).

### Offline Global Slow Oscillation Detection and Global Analysis of Stimulation Effects

To evaluate whether CLAS produced broader modifications of SO temporal organization beyond the local stimulation periods and account for limitations in online-detection procedure and offline detection cleaning, analyses were repeated using the complete set of offline-detected SOs identified throughout NREM sleep. This approach allowed the assessment of whether local stimulation effects translated into broader overnight modifications of slow oscillatory dynamics.

SOs (0.5-1 Hz) were automatically detected in artifact-free NREM (N2 and N3) epochs using YASA slow-waves detection function (Vallat & Walker, 2021), which is based on algorithms by Massimini et al., 2004 and Carrier et al., 2011. The frequency range parameter was set to YASA’s default (0.3-1.5 Hz) due to the 0.2 Hz transition band of its finite impulse response filter. The rest of the parameters used for SO detection were as follows: a negative deflection duration between 300 and 1500 ms (default value), a positive deflection duration between 100 and 1000 ms (default value), a negative peak amplitude below −40 µV with no upper limit, an unspecified positive peak amplitude, and a minimum peak-to-peak (PTP) amplitude threshold of 60 µV (Rosinvil et al., 2021; Lafrenière et al., 2023) with no upper limit. This reduced PTP amplitude criterion with respect to the traditional 75 µV threshold was set to account for the known reductions in SO amplitude associated with aging (Mander et al., 2017; Carrier et al., 2011; Rosinvil et al., 2021).

SO features (duration, frequency, PTP amplitude, positive slope, and negative and positive peak amplitudes) were obtained from the YASA slow-wave detection algorithm. Because YASA does not directly provide the negative slope, this parameter was calculated as the ratio between the negative peak amplitude and the time interval between the initial zero crossing and the negative peak. For each subject, feature values were averaged across all detected SOs, yielding a single value per recording.

### Sleep architecture and power spectral analysis

The total sleep time (TST), time spent in each sleep stage (N1, N2, N3, REM, and NREM [N2 + N3]), and time spent awake after sleep onset (WASO) were calculated for all subjects. Sleep onset was defined as the first occurrence of a N2 epoch. Sleep latency was calculated as the time elapsed between the start of the recording and sleep onset. The proportion of time spent in each stage relative to TST was determined, along with the percentages of N2 and N3 within total NREM sleep. N3 and REM sleep latencies from sleep onset were also measured. To further assess sleep fragmentation, in addition to WASO duration, the number of awakenings was quantified as the total count of transitions from NREM or REM sleep to W. Additionally, a sleep fragmentation index (SFI) was computed as the total number of awakenings and shifts to stage 1 (from NREM or REM sleep) divided by TST in hours. Certain recordings were fragmented into multiple files due to data acquisition interruptions caused by awakenings. The time elapsed between these files was accounted for in the calculation of total sleep time and awakenings.

In addition, a power spectral density analysis was performed for each night of each subject to assess the EEG power distribution across sleep stages and frequency rhythms of interest (SO: 0.5-1 Hz; delta: 1-4 Hz; theta: 4-8 Hz; alpha: 8-13 Hz; slow spindles: 9-12 Hz; fast spindles: 12-15 Hz; and beta: 15-30 Hz). The PSD was calculated using Welch’s method, implemented through the psd_array_welch function from the mne.time_frequency module. Artifact-free epochs were segmented into 10-second consecutive blocks with a 50% overlap. Each block was tapered by a single Hanning window before applying Fast Fourier Transformation that resulted in block power spectra with a frequency resolution of 0.1 Hz. Power spectra were then averaged across all blocks within each sleep stage. For each sleep stage, relative power in each frequency band was computed as the ratio between the area under the PSD curve within the corresponding frequency range and the total area under the PSD curve between 0.5 and 30 Hz.

### Statistics and Reproducibility

All statistical analyses were conducted in R (version 4.4.1) (R Core Team, 2024) and Python using SciPy (version 1.14.0) (Virtanen et al., 2020) and Pingouin (version 0.6.1) (Vallat, 2018). Plots were generated using Matplotlib (version 3.9.1) (Hunter, 2007).

Stimulation-related effects were evaluated by testing whether changes from the baseline to the experimental night differed between the STIM and SHAM groups. Descriptive changes were calculated within subjects as Δ = EXP - BASELINE and are reported as mean ± SEM for each group. Changes in percentages are expressed in percentage points. Statistical methods were selected according to the nature of each outcome, distinguishing conventional sleep measures, proportions representing two complementary categories, and compositions comprising multiple categories.

Sleep-stage durations, TST, WASO, sleep latencies, awakenings, SFI, relative spectral power, and SO morphological characteristics were analyzed using two-way mixed repeated-measures ANOVAs, with night as the within-subject factor and group as the between-subject factor.

Proportions representing two complementary categories were analyzed using beta mixed-effects regression with a logit link, implemented in glmmTMB (Brooks et al., 2017). Because the two proportions within each outcome sum to one, only one component was modeled. These outcomes comprised the proportion of N3 within NREM sleep, consecutive SOs among all detected SOs, train-start SOs among newly occurring SOs, and delta waves among validated online-detected events. For consecutive and train-start proportions, sensitivity analyses additionally included the corresponding recording-level SO density as a covariate. Separate models of consecutive SO proportions, train-start SO proportions, and SO density were fitted for the local morphology-cleaned, global offline-detected, and local non-morphology-cleaned datasets. Percentages were converted to proportions before modeling; zero values were replaced with 0.0001.

SO density was analyzed using linear mixed-effects models fitted by restricted maximum likelihood (REML). All beta and linear mixed-effects models included group, night, and their interaction as fixed effects and a subject-specific random intercept to account for repeated observations. Group and night were dummy-coded as SHAM = 0 and STIM = 1, and BASELINE = 0 and EXP = 1, respectively. Thus, the group coefficient represented the between-group difference during BASELINE, the night coefficient represented the EXP - BASELINE change within SHAM, and the interaction represented the between-group difference in change on the model’s respective scale. Fixed-effect inference used Wald z-tests for beta models and t-tests with Satterthwaite-approximated degrees of freedom for linear models.

To assess changes in the average number of SOs per train, within-subject changes in mean train length (EXP - BASELINE) were compared between STIM and SHAM using two-sided Welch’s t-tests.

Compositions comprising more than two categories were analyzed jointly using isometric log-ratio (ILR) transformations followed by change-score MANOVA. This approach was applied to sleep-stage composition and to train-length composition in each of the three SO datasets. For sleep-stage composition, Helmert-like partitions represented NREM versus REM, N1 versus NREM and REM, and wakefulness versus sleep. For train-length composition, an orthonormal Helmert basis provided one fewer ILR coordinate than the number of categories. Zero train-length proportions were replaced with 0.000001 before log-ratio transformation. Within-subject changes in the ILR coordinates (EXP - BASELINE) were entered jointly as the multivariate response, with group as the predictor. Between-group differences in compositional change were assessed using Pillai’s trace.

Exploratory component-specific analyses examined changes in the relative representation of individual train-length categories. Each category was compared with the geometric mean of all remaining train-length proportions using a normalized pivot ILR contrast. Subject-level changes in these contrasts were compared between groups using one-way ANOVAs. Holm correction was applied across all train-length categories within each dataset. These exploratory comparisons were performed irrespective of omnibus significance and interpreted as changes in relative composition rather than absolute changes in individual proportions.

Statistical significance was set at p < 0.05. Test statistics, degrees of freedom where applicable, and p-values are reported in the corresponding results sections and tables.

During manuscript preparation, the authors used ChatGPT (OpenAI; GPT-6.1 Terra) to refine English wording and improve clarity. All suggested revisions were reviewed and edited by the authors, who take full responsibility for the final content.

## Data availability

Data used in this study were collected by the Laboratorio de Sueño y Memoria at the Instituto Tecnológico de Buenos Aires (ITBA) and will be publicly available upon acceptance.

