## Supplementary Materials for "Closed-loop acoustic stimulation promotes local slow oscillation train initiation and persistence in elderly adults"

*Lucila Capurro, Iguazú 341, C1437 Ciudad Autónoma de Buenos Aires, Argentina

**Abstract**

Slow oscillations (SOs) during non-rapid eye movement (NREM) sleep support memory consolidation and metabolic waste clearance. Aging is associated with reduced SO amplitude and density and disrupted temporal organization, characterized by more isolated events and shorter SO trains. We investigated whether closed-loop acoustic stimulation (CLAS) could enhance the initiation and persistence of rhythmic SO activity in elderly adults. Twenty-seven participants underwent a baseline night followed by an experimental night, during which online-detected SOs were either stimulated using CLAS (STIM; n = 16) or left unstimulated (SHAM; n = 11). We quantified the proportions of isolated and consecutive SOs, train-initiation probability, and SO train length. Analyses assessed local effects around online-detected SOs and global effects across NREM sleep using offline detection. Locally, CLAS reduced the proportion of isolated SOs and increased the proportion of consecutive SOs, train-initiation probability, and mean train length. Global overnight effects were weaker but directionally consistent, primarily reflecting a shift toward longer SO trains. These changes occurred without a significant between-group difference in the change in SO density, and local effects remained significant after adjustment for SO density. Thus, CLAS promoted sustained rhythmic activity beyond changes in SO occurrence. These findings suggest that CLAS can modify the temporal organization of SOs in elderly adults, supporting its potential to mitigate age-related disruptions in SO dynamics. Temporal organization measures may complement conventional amplitude and density measures when evaluating CLAS effects. Further studies should establish whether these changes translate into cognitive benefits or improvements in other aspects of brain health.

**Temporal Distribution of Detections and Stimulations Across the Night**

Because slow-wave activity is not uniformly distributed across sleep and is more prominent during the first part of the night (Massimini et al., 2004), differences in the timing of detections or stimulation delivery could potentially bias comparisons between groups. Therefore, we examined the temporal distribution of online detections and delivered stimulations across the night.

For each subject, event times were normalized to the individual sleep period, defined from sleep onset to the end of sleep, and expressed as relative sleep time. Detection counts were then computed in ten equally spaced bins and normalized by the total number of detections for each subject. Mean distributions across subjects are shown in **Fig. S1**.

As expected, detections were concentrated toward the first half of the night, consistent with the higher prevalence of slow-wave activity early in sleep (**Fig. S1A,B**). Importantly, the temporal profiles of detections were highly similar between SHAM and STIM groups during both the BASELINE and EXP nights, indicating comparable sampling of sleep periods. Small shifts in the distribution of detections were observed across nights, with SHAM detections becoming slightly more concentrated in the first half of the night during the experimental session, whereas STIM detections exhibited a relatively longer tail toward later sleep periods. Importantly, these shifts would be expected to favor greater SO rhythmicity in SHAM and reduced rhythmicity in STIM, given the higher prevalence of organized SO activity early in the night. This pattern is opposite to the changes observed in the main analyses, indicating that differences in SO temporal organization cannot be explained by differences in the temporal distribution of detected events.

To verify that auditory stimulation followed the same temporal pattern as detected events, we compared the distributions of detections and delivered stimulations during the experimental night in the STIM group (**Fig. S1C**). Both distributions almost completely overlapped, demonstrating that stimulation delivery closely tracked the occurrence of detected events. A minor divergence was observed only towards the end of the night because a few detections were generated after stimulation had been manually discontinued by the supervising technicians in some subjects (see **Fig. S2**)

Overall, these analyses indicate that detections and stimulations were similarly distributed across the course of the night in all experimental conditions. Therefore, the differences reported in the main analyses cannot be explained by differences in the temporal occurrence of detected events.

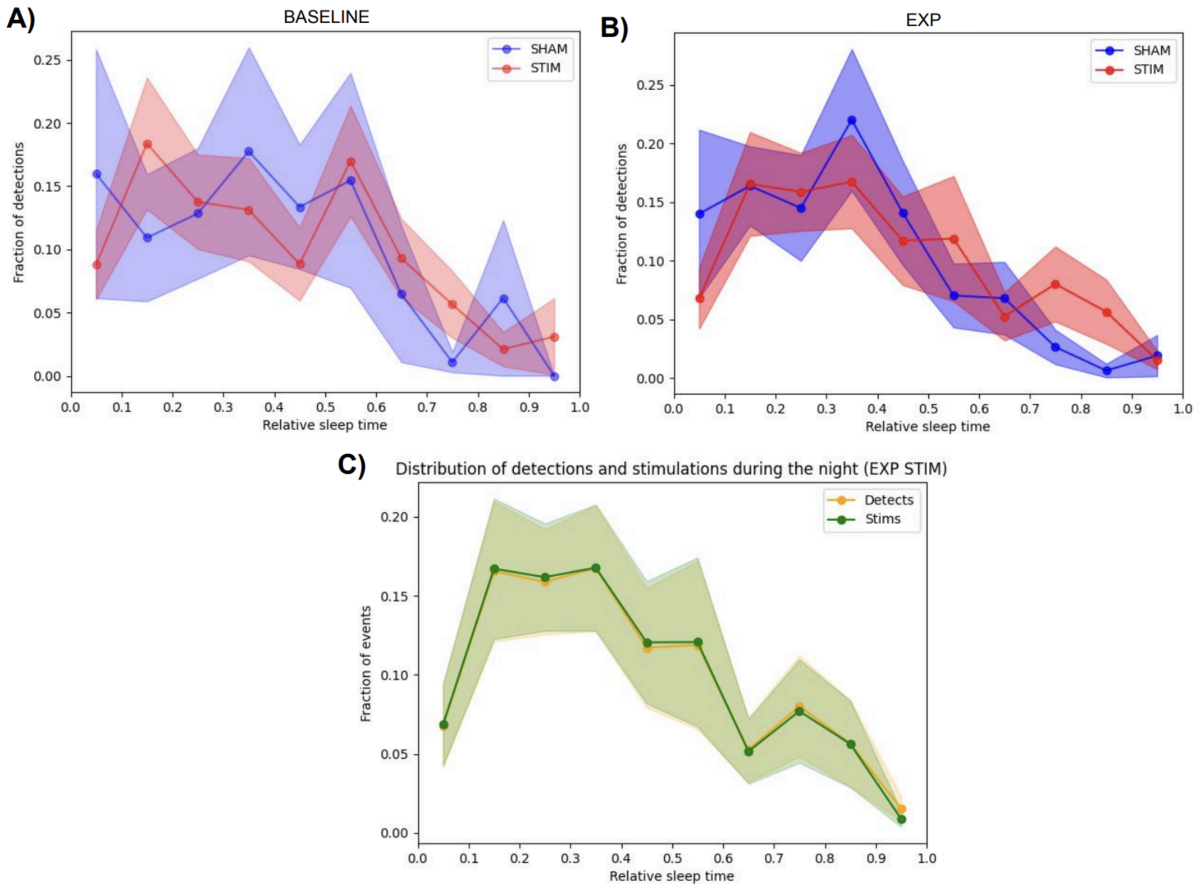

**Figure S1**. **Temporal distribution of online detections and stimulations across the night.** Event times were expressed as relative sleep time (from sleep onset to sleep end) and divided into ten equal intervals. Values represent the mean fraction of events in each interval across subjects, and shaded areas indicate SEM. **(A)** Distribution of online detections during the BASELINE night for SHAM (blue, N = 11) and STIM (red, N = 16) groups. **(B)** Distribution of online detections during the EXP night. **(C)** Comparison between the distributions of detections (orange) and delivered auditory stimulations (green) during the experimental night in the STIM group.

**Online Slow-Wave Detection Cleaning and Event Retention**

The local analyses described in the main text were performed on a subset of online detections that remained after several preprocessing steps designed to remove events that could not be reliably interpreted as slow waves. **Figure S2** summarizes the proportion of detections excluded at each stage of the preprocessing pipeline. Percentages for each intermediate step were calculated relative to the detections surviving the previous preprocessing stage.

First, detections occurring during interpolated periods were removed. Interpolated periods corresponded to epochs containing interpolated samples, as described in the Offline Preprocessing section. This step had a negligible impact on the dataset, with nearly all subjects showing rejection rates below 1% and only a few isolated cases exceeding 5%.

Second, for the STIM experimental night, detections that were not followed by auditory stimulation were excluded. Because detection and stimulation could be enabled or disabled independently by the supervising technicians, a small number of detections were generated after stimulation delivery had been paused, primarily toward the end of stimulation periods. The proportion of such detections was generally low, with a median rejection rate below 1%.

Third, detections were restricted to offline-scored N2 and N3 sleep. This step removed a modest proportion of events in most subjects, typically between 1% and 5% of the detections surviving the previous preprocessing stages. However, two cases exhibited substantially higher rejection rates, reaching approximately 20% and 42%.

Finally, the waveform reconstruction procedure described in the main text was applied to identify detections corresponding to well-defined slow waves. Events for which the negative peak or the associated waveform landmarks could not be reliably identified were classified as erroneous online detections and excluded. For most subjects, this step resulted in minimal additional event loss. Notably, the case showing the largest rejection rates during NREM filtering also exhibited elevated rejection rates at this stage.

Across all preprocessing stages, the median proportion of discarded detections remained below 5% in all groups and recording nights. The cumulative proportion of excluded detections was below 10% for the vast majority of subjects. Nevertheless, a small number of cases exhibited higher overall rejection rates (approximately 10-27%), and one case showed a markedly elevated rejection rate of 72%, primarily due to detections occurring outside N2/N3 sleep and the subsequent failure to satisfy the slow-wave morphology criteria (**Fig. S2**).

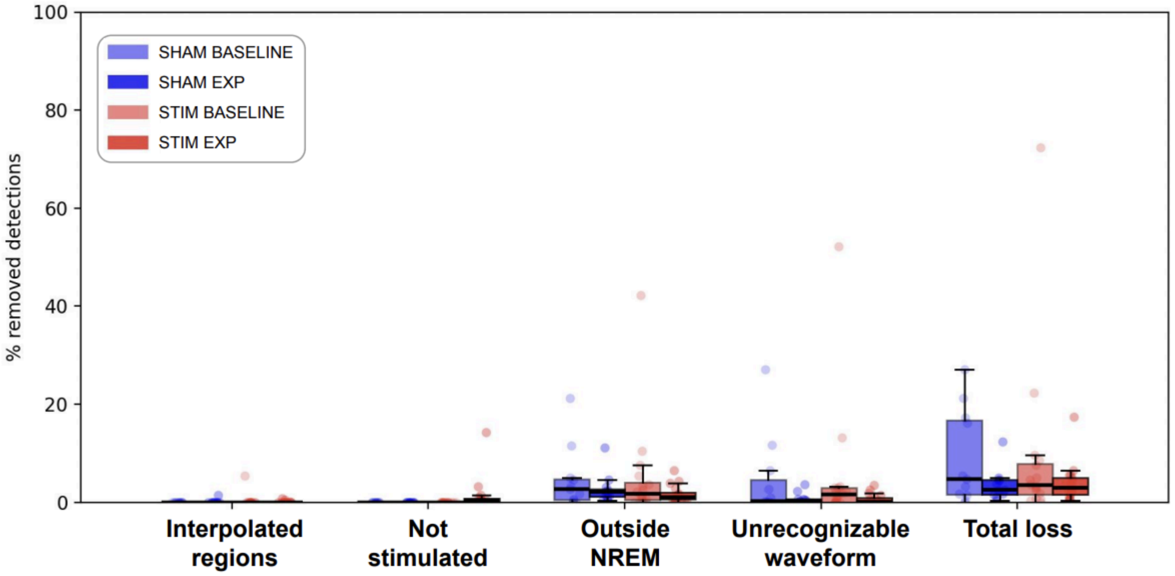

**Figure S2**. **Detection losses across preprocessing stages.** Distribution of the percentage of online detections excluded at each preprocessing step. "Interpolated Regions" indicates detections occurring during artifact periods containing interpolated samples. "Not stimulated" corresponds to detections from the STIM experimental night that were not followed by auditory stimulation. "Outside NREM sleep" indicates detections occurring outside offline-scored N2 or N3 sleep. "Unrecognizable waveform" corresponds to events that failed the offline waveform reconstruction procedure and therefore did not meet the slow-wave morphology criteria. Percentages for each intermediate step were computed relative to the detections surviving the previous preprocessing stage, whereas "Total loss" represents the cumulative proportion of detections discarded relative to the original number of online raw detections. Individual points represent subjects and boxplots summarize group distributions.

### Statistical results for local and global SO temporal organization

**Tables S1** and **S2** provide statistical results supporting the local and global SO temporal organization analyses presented in the main manuscript **(Figs. 2 and 3**, respectively). These include group, night, and group x night coefficients for consecutive SO proportions, train-start SO proportions, and SO density, together with SO-density covariate effects in the adjusted proportion models. The tables also report between-group comparisons of changes in mean train length, omnibus ILR-MANOVAs of train-length composition, and exploratory pivot ILR contrasts comparing each train-length proportion with the geometric mean of the remaining proportions, with Holm correction within each dataset. Corresponding results for local detections retained without waveform morphology rejection are provided below in **Table S3** and described in “Robustness of Results to Detection Preprocessing”.

**Table S1. Local SO temporal organization after waveform morphology filtering.**

| **A. Proportion models without SO density adjustment** | | | | |
|---|---|---|---|---|
| Outcome | Group (STIM) | Night (EXP) | Interaction | |
| Consecutive SOs | b = -0.688, z = -1.685, p = 0.092 | b = 0.108, z = 0.305, p = 0.761 | b = 1.209, z = 2.555, **p = 0.011** | |
| Train-start SOs | b = -0.648, z = -1.729, p = 0.084 | b = 0.105, z = 0.325, p = 0.746 | b = 1.093, z = 2.536, **p = 0.011** | |
| **B. Proportion models adjusted for SO density** | | | | |
| Outcome | Group (STIM) | Night (EXP) | Interaction | SO density |
| Consecutive SOs | b = -0.164, z = -0.440, p = 0.660 | b = 0.325, z = 1.011, p = 0.312 | b = 0.876, z = 2.077, **p = 0.038** | b = 0.297, z = 5.076, **p < 0.001** |

| | | | | |
|---|---|---|---|---|
| Train-start SOs | b = -0.155, z = -0.449, p = 0.653 | b = 0.326, z = 1.102, p = 0.270 | b = 0.775, z = 1.975, **p = 0.048** | b = 0.247, z = 5.132, **p < 0.001** |
| **C. SO density** | | | | |
| Outcome | Group (STIM) | Night (EXP) | Interaction | |
| NREM SO density | b = -1.785, t(39.79) = -2.084, **p = 0.044** | b = -0.739, t(25.00) = -1.129, p = 0.270 | b = 1.259, t(25.00) = 1.479, p = 0.152 | |
| **D. Mean train length and train-length composition** | | | | |
| Outcome | Estimate | Test statistic | p | |
| Mean train length | ΔSTIM - ΔSHAM = 0.254 SOs/train; 95% CI [0.062, 0.446] | t(24.34) = 2.727 | **0.012** | |
| Train-length composition | Pillai's trace = 0.334 | F(5,21) = 2.109 | 0.104 | |
| **E. Exploratory component-specific pivot ILR contrasts** | | | | |
| Train length | ΔSTIM - ΔSHAM (ILR units) | F(1,25) | p | Holm-adjusted p |
| 2 | -3.644 | 7.294 | **0.012** | 0.073 |
| 3 | 2.458 | 1.330 | 0.260 | 1.000 |
| 4 | 4.458 | 6.593 | **0.017** | 0.083 |
| 5 | -0.327 | 0.027 | 0.871 | 1.000 |
| 6 | -1.856 | 1.176 | 0.289 | 1.000 |
| 7 | -1.089 | 0.689 | 0.414 | 1.000 |

*Statistical results corresponding to Fig. 2. Panels A and B report regression coefficients (b), Wald z-statistics, and p-values from beta mixed-effects models of consecutive SOs among all detected SOs and train-start SOs among newly occurring SOs. Coefficients are expressed on the logit scale. Panel B additionally includes SO density as a covariate. Panel C reports coefficients on the SO-density scale, t-statistics with Satterthwaite-approximated degrees of freedom in parentheses, and p-values from a linear mixed-effects model. All mixed-effects models included a subject-specific random intercept. SHAM and BASELINE were the reference categories: the group coefficient represents the STIM - SHAM difference during BASELINE, the night coefficient represents the EXP - BASELINE change within SHAM, and the group x night coefficient represents the between-group difference in that change. In density-adjusted models, these contrasts are conditional on SO density. Panel D reports between-group differences in subject-level changes (Δ = EXP - BASELINE) in mean train length, evaluated using two-sided Welch's t-tests, and omnibus change-score MANOVAs of ILR-transformed train-length compositions, evaluated using Pillai's trace. Mean train length was calculated as the proportion-weighted mean number of SOs per train. Confidence intervals are pointwise 95% intervals. Panel E reports exploratory pivot ILR contrasts comparing each train-length proportion with the geometric mean of all remaining train-length proportions. Subject-level changes in these contrasts were compared between groups using one-way ANOVAs. Holm correction was applied across all train-length contrasts within each dataset. Positive differences indicate greater EXP - BASELINE changes in the corresponding log-ratio in STIM than SHAM. Significant p-values are highlighted in bold.*

**Table S2. Global overnight SO temporal organization.**

| **A. Proportion models without SO density adjustment** | | | |
|---|---|---|---|
| Outcome | Group (STIM) | Night (EXP) | Interaction |

| | | | | |
|---|---|---|---|---|
| Consecutive SOs | b = 0.034, z = 0.155, p = 0.877 | b = 0.226, z = 1.452, p = 0.146 | b = 0.109, z = 0.538, p = 0.590 | |
| Train-start SOs | b = 0.095, z = 0.531, p = 0.595 | b = 0.236, z = 1.857, p = 0.063 | b = -0.006, z = -0.038, p = 0.970 | |
| **B. Proportion models adjusted for SO density** | | | | |
| Outcome | Group (STIM) | Night (EXP) | Interaction | SO density |
| Consecutive SOs | b = 0.095, z = 0.635, p = 0.525 | b = 0.052, z = 0.340, p = 0.734 | b = 0.222, z = 1.127, p = 0.260 | b = 0.363, z = 7.007, **p < 0.001** |
| Train-start SOs | b = 0.151, z = 1.182, p = 0.237 | b = 0.099, z = 0.766, p = 0.444 | b = 0.069, z = 0.419, p = 0.676 | b = 0.276, z = 6.854, **p < 0.001** |
| **C. SO density** | | | | |
| Outcome | Group (STIM) | Night (EXP) | Interaction | |
| NREM SO density | b = -0.180, t(27.50) = -0.393, p = 0.697 | b = 0.494, t(25.00) = 3.219, **p = 0.004** | b = -0.311, t(25.00) = -1.560, p = 0.131 | |
| **D. Mean train length and train-length composition** | | | | |
| Outcome | Estimate | Test statistic | p | |
| Mean train length | ΔSTIM - ΔSHAM = 0.266 SOs/train; 95% CI [0.008, 0.525] | t(21.46) = 2.138 | **0.044** | |
| Train-length composition | Pillai's trace = 0.185 | F(7,19) = 0.616 | 0.736 | |
| **E. Exploratory component-specific pivot ILR contrasts** | | | | |
| Train length | ΔSTIM - ΔSHAM (ILR units) | F(1,25) | p | Holm-adjusted p |
| 2 | -1.430 | 1.596 | 0.218 | 1.000 |
| 3 | -0.813 | 0.646 | 0.429 | 1.000 |
| 4 | 0.370 | 0.053 | 0.820 | 1.000 |
| 5 | 2.608 | 2.041 | 0.166 | 1.000 |
| 6 | 1.355 | 0.413 | 0.526 | 1.000 |
| 7 | 0.265 | 0.025 | 0.876 | 1.000 |
| 8 | -0.984 | 0.364 | 0.552 | 1.000 |
| 9 | -1.372 | 0.575 | 0.455 | 1.000 |

*Statistical results corresponding to Fig. 3. Panels A and B report regression coefficients (b), Wald z-statistics, and p-values from beta mixed-effects models of consecutive SOs among all detected SOs and train-start SOs among newly occurring SOs. Coefficients are expressed on the logit scale. Panel B additionally includes SO density as a covariate. Panel C reports coefficients on the SO-density scale, t-statistics with Satterthwaite-approximated degrees of freedom in parentheses, and p-values from a linear mixed-effects model. All mixed-effects models included a subject-specific random intercept. SHAM and BASELINE were the reference categories: the group coefficient represents the STIM - SHAM difference during BASELINE, the night coefficient represents the EXP - BASELINE change within SHAM, and the group x night coefficient represents the between-group difference in that change. In density-adjusted models, these contrasts are conditional on SO density. Panel D reports between-group differences in subject-level changes (Δ = EXP - BASELINE) in mean train length, evaluated using two-sided Welch's t-tests, and omnibus change-score MANOVAs of ILR-transformed train-length compositions, evaluated using Pillai's trace. Mean train length was calculated as the proportion-weighted mean number of SOs per train. Confidence intervals are pointwise 95% intervals. Panel E reports exploratory pivot ILR contrasts comparing each train-length proportion with the geometric mean of all remaining train-length proportions. Subject-level changes in these contrasts were compared between groups using one-way ANOVAs. Holm correction was applied across all train-length contrasts within each dataset. Positive differences indicate greater EXP - BASELINE changes in the corresponding log-ratio in STIM than SHAM. Significant p-values are highlighted in bold. Significant p-values are highlighted in bold.*

**Robustness of Results to Detection Preprocessing**

To assess whether the local stimulation effects reported in the main manuscript depended on the waveform-based rejection procedure, the local analyses were repeated using all NREM detections retained after artifact and stage filtering, regardless of waveform morphology.

Consistent with the morphology-cleaned dataset shown in **Figure 2**, STIM showed a greater increase in the proportion of consecutive SOs and a complementary decrease in isolated SOs from BASELINE to EXP compared with SHAM (group x night interaction: $b = 2.402$, $z = 4.190$, $p < 0.001$; **Fig. S3A**). Among newly occurring SOs, STIM also showed a greater increase in the probability of initiating a train rather than remaining isolated ($b = 2.177$, $z = 4.122$, $p < 0.001$; **Fig. S3B**). Both effects remained significant after accounting for SO density (consecutive SOs: $b = 2.142$, $z = 3.839$, $p < 0.001$; train-start SOs: $b = 1.923$, $z = 3.693$, $p < 0.001$).

This greater propensity to form trains was accompanied by a greater increase in mean train length in STIM than in SHAM (ΔSTIM = 0.179 ± 0.067 vs ΔSHAM = -0.075 ± 0.076 SOs per train; Welch's $t(22.59) = 2.495$, $p = 0.020$; **Fig. S3C**). Consistent with this finding, the train-length distribution showed a descriptive shift toward longer trains in STIM relative to SHAM (**Fig. S3D**). The joint ILR-MANOVA of the complete train-length composition narrowly missed the prespecified significance threshold (Pillai's trace = 0.390, $F(5,21) = 2.684$, $p = 0.05003$).

Exploratory component-specific comparisons indicated a greater increase in the log-ratio of four-wave trains to the geometric mean of the remaining train-length proportions in STIM than in SHAM ($F(1,25) = 8.360$, $p = 0.008$, Holm-adjusted $p = 0.047$). The corresponding relative decrease in two-wave trains reached nominal significance but did not survive correction ($F(1,25) = 7.348$, $p = 0.012$, adjusted $p = 0.060$). No other comparison reached significance. Because the omnibus test did not meet the significance threshold, these component-specific findings were interpreted as exploratory.

These changes occurred without a statistically significant between-group difference in the change in NREM SO density (group x night interaction: $b = 1.457$, $t(25) = 1.607$, $p = 0.121$; **Fig. S3E**). Complete statistical results are provided in **Table S3**. Together, these findings show that the greater participation of SOs in trains, increased probability of train initiation, and greater mean train length observed locally were also present without waveform-based rejection, supporting their robustness to this preprocessing step.

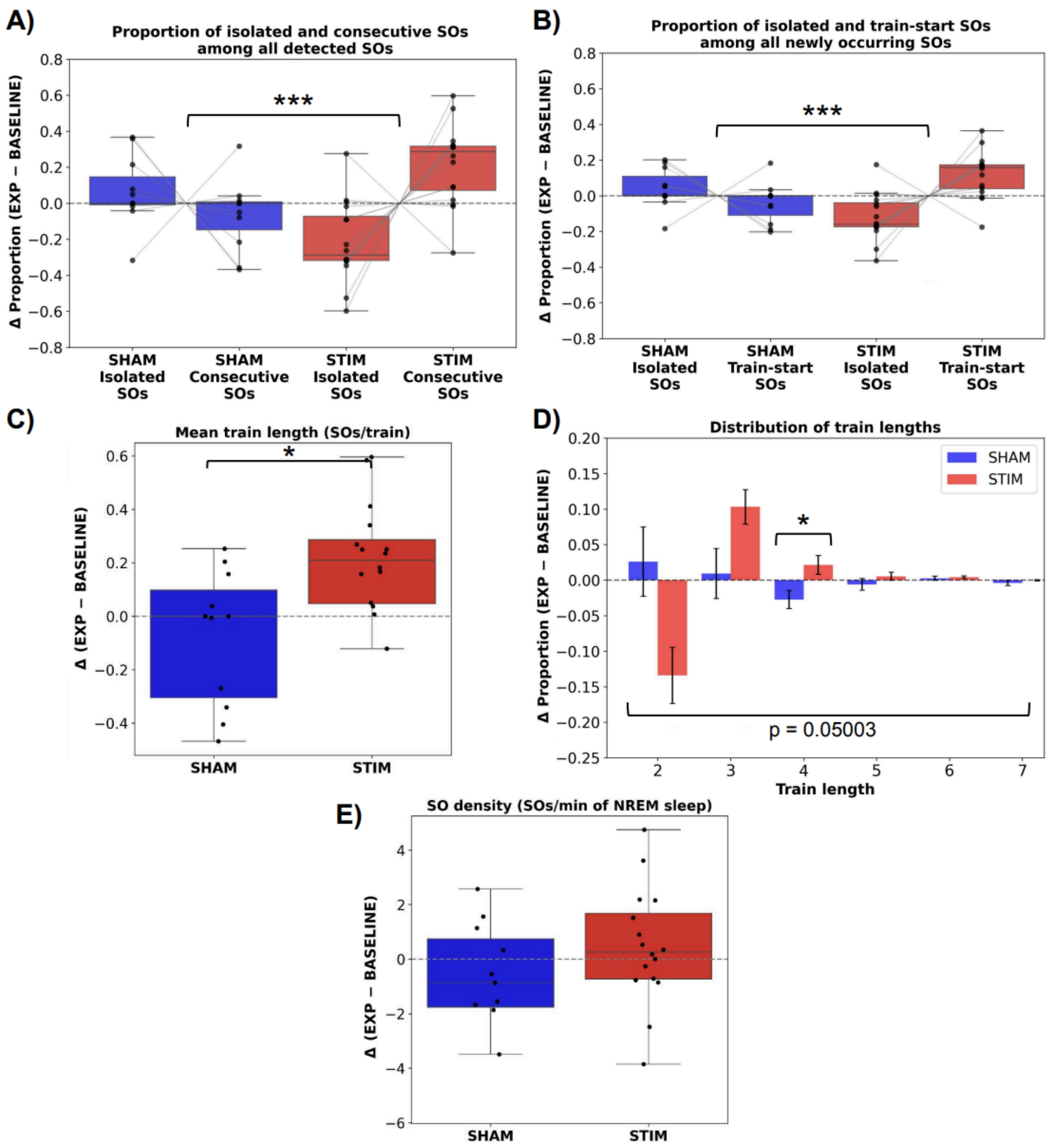

**Figure S3. Robustness of local CLAS effects to waveform-based preprocessing.** Within-subject changes between the experimental and baseline nights (EXP - BASELINE) are shown for SHAM (blue, n = 11) and STIM (red, n = 16). **(A)** Proportions of isolated and consecutive SOs among all detected SOs. **(B)** Proportions of isolated and train-start SOs among newly occurring SOs. **(C)** Mean train length, calculated as the proportion-weighted mean number of SOs per train. **(D)** Changes in train-length proportions, shown as group means ± SEM. **(E)** Changes in NREM SO density. Positive values indicate increases from BASELINE to EXP. Individual dots represent subjects. ***p < 0.001; **p < 0.01; *p < 0.05.

**Table S3. Local SO temporal organization without waveform morphology rejection.**

| **A. Proportion models without SO density adjustment** | | | |
|---|---|---|---|
| Outcome | Group (STIM) | Night (EXP) | Interaction |
| Consecutive SOs | b = -1.414, z = -2.815, **p = 0.005** | b = -0.372, z = -0.890, p = 0.374 | b = 2.402, z = 4.190, **p < 0.001** |

| | | | | |
|---|---|---|---|---|
| Train-start SOs | b = -1.322, z = -2.881, **p = 0.004** | b = -0.342, z = -0.888, p = 0.374 | b = 2.177, z = 4.122, **p < 0.001** | |
| **B. Proportion models adjusted for SO density** | | | | |
| Outcome | Group (STIM) | Night (EXP) | Interaction | SO density |
| Consecutive SOs | b = -0.958, z = -1.919, p = 0.055 | b = -0.184, z = -0.444, p = 0.657 | b = 2.142, z = 3.839, **p < 0.001** | b = 0.259, z = 3.311, **p < 0.001** |
| Train-start SOs | b = -0.887, z = -1.942, p = 0.052 | b = -0.149, z = -0.390, p = 0.696 | b = 1.923, z = 3.693, **p < 0.001** | b = 0.225, z = 3.481, **p < 0.001** |
| **C. SO density** | | | | |
| Outcome | Group (STIM) | Night (EXP) | Interaction | |
| NREM SO density | b = -1.973, t(40.32) = -2.198, **p = 0.034** | b = -1.004, t(25.00) = -1.439, p = 0.163 | b = 1.457, t(25.00) = 1.607, p = 0.121 | |
| **D. Mean train length and train-length composition** | | | | |
| Outcome | Estimate | Test statistic | p | |
| Mean train length | ΔSTIM - ΔSHAM = 0.254 SOs/train; 95% CI [0.043, 0.465] | t(22.59) = 2.495 | **0.020** | |
| Train-length composition | Pillai's trace = 0.390 | F(5,21) = 2.684 | **0.05003** | |
| **E. Exploratory component-specific pivot ILR contrasts** | | | | |
| Train length | ΔSTIM - ΔSHAM (ILR units) | F(1,25) | p | Holm-adjusted p |
| 2 | -3.727 | 7.348 | **0.012** | 0.060 |
| 3 | 2.335 | 1.228 | 0.278 | 1.000 |
| 4 | 5.071 | 8.360 | **0.008** | **0.047** |
| 5 | -0.565 | 0.084 | 0.775 | 1.000 |
| 6 | -1.872 | 1.150 | 0.294 | 1.000 |
| 7 | -1.241 | 0.812 | 0.376 | 1.000 |

*Statistical results corresponding to Fig. S3. Sections A and B report regression coefficients (b), Wald z-statistics, and p-values from beta mixed-effects models of consecutive SOs among all detected SOs and train-start SOs among newly occurring SOs. Coefficients are expressed on the logit scale. Panel B additionally includes SO density as a covariate. Panel C reports coefficients on the SO-density scale, t-statistics with Satterthwaite-approximated degrees of freedom in parentheses, and p-values from a linear mixed-effects model. All mixed-effects models included a subject-specific random intercept. SHAM and BASELINE were the reference categories: the group coefficient represents the STIM - SHAM difference during BASELINE, the night coefficient represents the EXP - BASELINE change within SHAM, and the group x night coefficient represents the between-group difference in that change. In density-adjusted models, these contrasts are conditional on SO density. Panel D reports between-group differences in subject-level changes (Δ = EXP - BASELINE) in mean train length, evaluated using two-sided Welch's t-tests, and omnibus change-score MANOVAs of ILR-transformed train-length compositions, evaluated using Pillai's trace. Mean train length was calculated as the proportion-weighted mean number of SOs per train. Confidence intervals are pointwise 95% intervals. Panel E reports exploratory pivot ILR contrasts comparing each train-length proportion with the geometric mean of all remaining train-length proportions. Subject-level changes in these contrasts were compared between groups using one-way ANOVAs. Holm correction was applied across all train-length contrasts within each dataset. Positive differences indicate greater EXP*

*- BASELINE changes in the corresponding log-ratio in STIM than SHAM. Significant p-values are highlighted in bold. Significant p-values are highlighted in bold.*

**Morphological Characteristics of Valid Online-Detected Slow Oscillations**

Because the classification of SOs into isolated and consecutive events depends on the temporal spacing between neighboring detected waves, we examined whether the effects reported in the main manuscript could be explained by systematic differences in the characteristics of the detected waves themselves rather than by genuine changes in their temporal organization. To this end, conventional morphological features were computed for the locally detected SOs that survived the preprocessing pipeline. The analyzed parameters included peak-to-peak (PTP) amplitude, negative and positive peak amplitudes, slopes, and frequency.

No significant group effects or group x night interactions were observed for any morphological parameter (**Table S4**), indicating that CLAS did not induce systematic changes in the shape of locally detected SOs. Only positive peak amplitude and frequency exhibited significant main effects of night, reflecting changes that occurred similarly in both SHAM and STIM groups. In particular, SO frequency decreased during the experimental night, corresponding to a longer oscillatory period and longer SO duration. Importantly, this effect would be expected to increase the intervals between successive SOs and thereby reduce, rather than increase, the probability of observing consecutive events. Thus, the enhanced SO train persistence observed in the main analyses cannot be explained by a shift toward faster oscillations, by differences in waveform morphology, or by systematic changes in the detection properties of the waves themselves. Instead, the increased prevalence of consecutive SOs and longer SO trains reflects a genuine reorganization of SO temporal dynamics rather than changes in the morphology or detectability of individual waves.

**Table S4. Morphological characteristics of locally detected SOs.**

| Variable | ΔSHAM | ΔSTIM | Night | Group | Interaction |
|---|---|---|---|---|---|
| **Negative peak (uV)** | -5.32 ± 3.61 | -1.37 ± 1.73 | F(1,25)=2.78, p=0.108 | F(1,25)=0.07, p=0.794 | F(1,25)=1.18, p=0.287 |
| **Positive peak (uV)** | 2.01 ± 0.94 | 1.36 ± 0.71 | F(1,25)=8.14, **p=0.009** | F(1,25)=0.23, p=0.634 | F(1,25)=0.32, p=0.576 |
| **PTP amplitude (uV)** | 7.33 ± 4.46 | 2.72 ± 2.38 | F(1,25)=4.04, p=0.055 | F(1,25)=0.13, p=0.725 | F(1,25)=0.98, p=0.332 |
| **Negative slope (uV/s)** | -31.15 ± 21.18 | -3.55 ± 10.61 | F(1,25)=1.93, p=0.177 | F(1,25)=0.09, p=0.772 | F(1,25)=1.62, p=0.214 |
| **Positive slope (uV/s)** | 13.74 ± 11.18 | -7.64 ± 5.54 | F(1,25)=0.04, p=0.850 | F(1,25)=0.32, p=0.574 | F(1,25)=3.53, p=0.072 |
| **Frequency (Hz)** | -0.03 ± 0.02 | -0.06 ± 0.02 | F(1,25)=7.41, **p=0.012** | F(1,25)=0.54, p=0.467 | F(1,25)=0.89, p=0.356 |

*Values are expressed as the mean ± SEM of the change between the experimental and baseline nights (Δ = EXP - BASELINE) for the SHAM (N = 11) and STIM (N = 16) groups. Statistical analyses were performed using two-way mixed repeated-measures ANOVAs with night (BASELINE, EXP) as the within-subject factor and group (SHAM, STIM) as the between-subject factor. P values for the main effects of night and group and for the group x night interaction are shown. Significant p-values are highlighted in bold.*

To complement the quantitative analysis of morphological features, average SO waveforms were computed for each group and recording night (**Fig. S4**). Mean waveforms showed highly similar shapes across groups, consistent with the absence of significant group effects or group x night interactions in any morphological parameter.

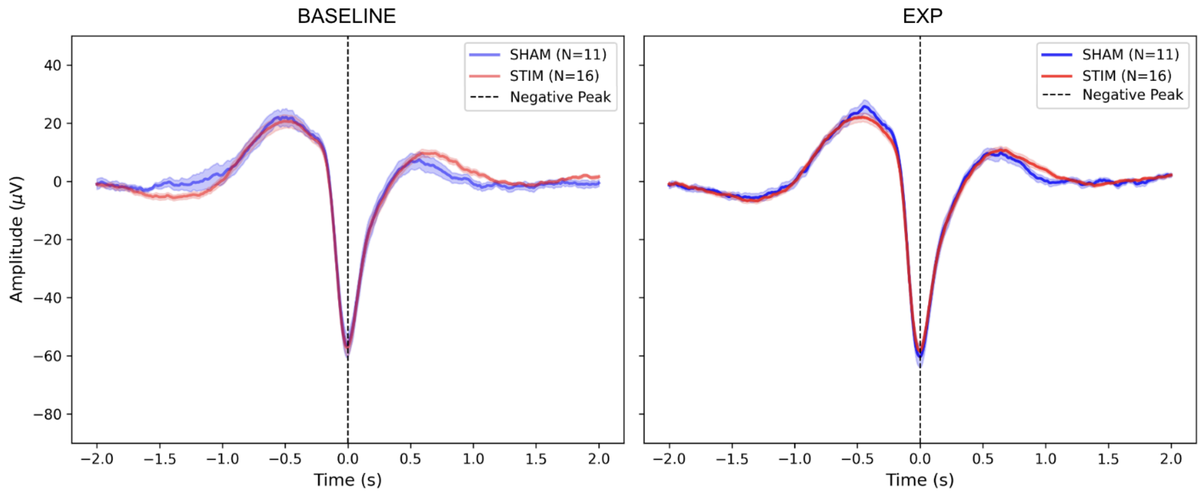

**Figure S4. Average local slow-wave morphology across groups and nights.** Grand-average waveforms of the valid online-detected slow waves centered on the negative peak (0 s) are shown for the SHAM (blue, N = 11) and STIM (red, N = 16) groups during the baseline (left) and experimental (EXP, right) nights. Average waveforms were first computed at the subject level after randomly subsampling SOs to match the number of events between BASELINE and EXP within each subject, and group averages were then obtained across subjects.

**Frequency Distribution of Valid Online-Detected Events**

Because the online detection algorithm relied on a broad low-pass filter and was optimized for real-time stimulation rather than strict waveform classification, validated events were not necessarily restricted to canonical slow oscillations (≤1 Hz) and could also include faster delta waves. We therefore examined the frequency distribution of all validated online-detected events to determine whether differences in SO temporal organization could be explained by changes in the types of waves detected.

Frequency histograms revealed highly similar distributions across groups and nights (**Fig. S5**). In all conditions, the distributions were strongly skewed toward frequencies below 1 Hz, indicating that most validated events corresponded to slow oscillations. Nevertheless, a progressively decreasing tail extended toward higher frequencies, reaching approximately 2.5 Hz, reflecting the presence of a smaller proportion of faster delta waves. This frequency estimate was derived from the waveform reconstruction procedure described in the Methods section rather than from the online detection filter itself.

Importantly, no evidence was found for a shift toward higher frequencies in the STIM group during the experimental night. On the contrary, the average event frequency showed a small decrease that occurred similarly in both groups (**Table S4**), corresponding to slightly longer oscillatory periods. Thus, the enhanced SO train initiation and persistence observed in the main analyses cannot be attributed to an increased contribution of faster delta waves or to systematic differences in the frequency composition of the detected events. Instead, the predominance of frequencies below 1 Hz across all conditions indicates that the observed effects primarily reflect changes in the temporal organization of slow oscillatory activity.

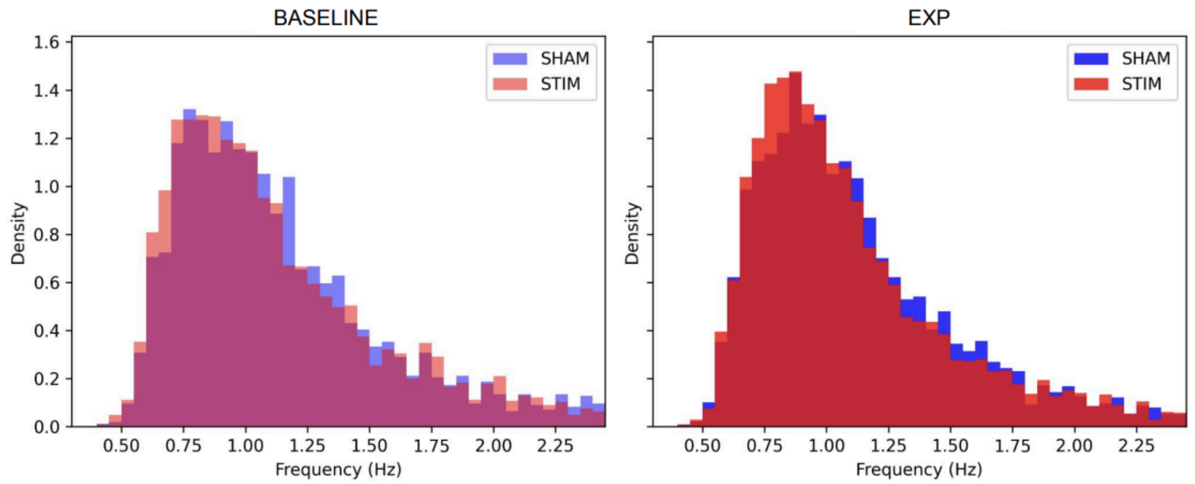

**Figure S5. Frequency distribution of valid online-detected events.** Frequency histograms of all validated online detections after artifact rejection, NREM filtering, and waveform reconstruction are shown for the SHAM (blue) and STIM (red) groups during the baseline (left) and experimental (EXP, right) nights. Event frequency was estimated from the reconstructed waveform as the inverse of the interval between the initial and final zero crossings. Histograms are shown as probability densities using 0.05-Hz bins.

To further quantify the frequency composition of detected events, reconstructed waves were classified as SOs (≤1 Hz) or delta waves (>1 Hz). Descriptively, the proportion of SOs increased from BASELINE to EXP by 3.16 ± 2.21 % in SHAM and 3.11 ± 1.62 % in STIM, with complementary decreases in delta-wave proportions. However, the beta mixed-effects analysis did not provide evidence of differential change between groups in the proportion of delta waves (group × night interaction: b = -0.004, z = -0.035, p = 0.972). Neither the between-group difference during BASELINE (b = -0.305, z = -1.236, p = 0.216) nor the BASELINE-to-EXP change within SHAM (b = -0.135, z = -1.563, p = 0.118) was statistically significant. Thus, although the experimental night was associated with a modest shift toward a greater proportion of SOs, this shift occurred similarly in both groups and therefore cannot account for the stimulation-related increases in SO train initiation and persistence.

**Morphological Characteristics of Offline-Detected Slow Oscillations**

To determine whether the CLAS-induced increase in SO train persistence could be explained by systematic differences in the characteristics of the detected oscillations, we compared conventional morphological features of all offline-detected NREM SOs identified with the YASA algorithm. For each recording, mean values of negative and positive peak amplitudes, peak-to-peak (PTP) amplitude, positive and negative slopes, and frequency were computed across all detected events, and changes between the EXP and BASELINE nights were compared between groups (**Table S5**).

No significant main effects of group or group x night interactions were observed for any morphological parameter. Likewise, no significant night effects were detected. Changes in all waveform characteristics were small and comparable between SHAM and STIM subjects. These findings indicate that the greater SO train persistence observed in the main analyses cannot be attributed to systematic differences in the morphology of the oscillations or to changes in the criteria used by the offline detection algorithm.

**Table S5. Morphological characteristics of offline-detected SOs.**

| Variable | ΔSHAM | ΔSTIM | Night | Group | Interaction |
|---|---|---|---|---|---|
| **Negative peak (uV)** | 0.34 ± 1.37 | -0.76 ± 0.41 | F(1,25)=0.26, p=0.615 | F(1,25)=1.50, p=0.232 | F(1,25)=0.79, p=0.382 |
| **Positive peak (uV)** | 0.33 ± 1.18 | 0.43 ± 0.58 | F(1,25)=0.43, p=0.516 | F(1,25)=0.02, p=0.885 | F(1,25)=0.01, p=0.934 |

| | | | | | |
|---|---|---|---|---|---|
| **PTP amplitude (uV)** | -0.01 ± 2.28 | 1.19 ± 0.80 | F(1,25)=0.45, p=0.508 | F(1,25)=0.20, p=0.655 | F(1,25)=0.32, p=0.575 |
| **Negative slope (uV/s)** | 0.92 ± 4.08 | -0.90 ± 2.19 | F(1,25)=0.01, p=0.941 | F(1,25)=0.37, p=0.550 | F(1,25)=0.18, p=0.674 |
| **Positive slope (uV/s)** | -1.60 ± 8.51 | 0.68 ± 4.15 | F(1,25)=0.00, p=0.953 | F(1,25)=0.15, p=0.698 | F(1,25)=0.07, p=0.794 |
| **Frequency (Hz)** | -0.01 ± 0.01 | -0.00 ± 0.01 | F(1,25)=0.45, p=0.508 | F(1,25)=0.46, p=0.505 | F(1,25)=0.17, p=0.683 |

*Values are expressed as the mean ± SEM of the change between the experimental and baseline nights (Δ = EXP - BASELINE) for the SHAM (N = 11) and STIM (N = 16) groups. Statistical analyses were performed using two-way mixed repeated-measures ANOVAs with night (BASELINE, EXP) as the within-subject factor and group (SHAM, STIM) as the between-subject factor. P values for the main effects of night and group and for the group x night interaction are shown.*

Again, to complement the quantitative analysis of morphological features, average SO waveforms were computed for each group and recording night (**Fig. S6**). Mean waveforms showed highly similar shapes across nights and groups, consistent with the absence of significant group effects or group x night interactions in any morphological parameter.

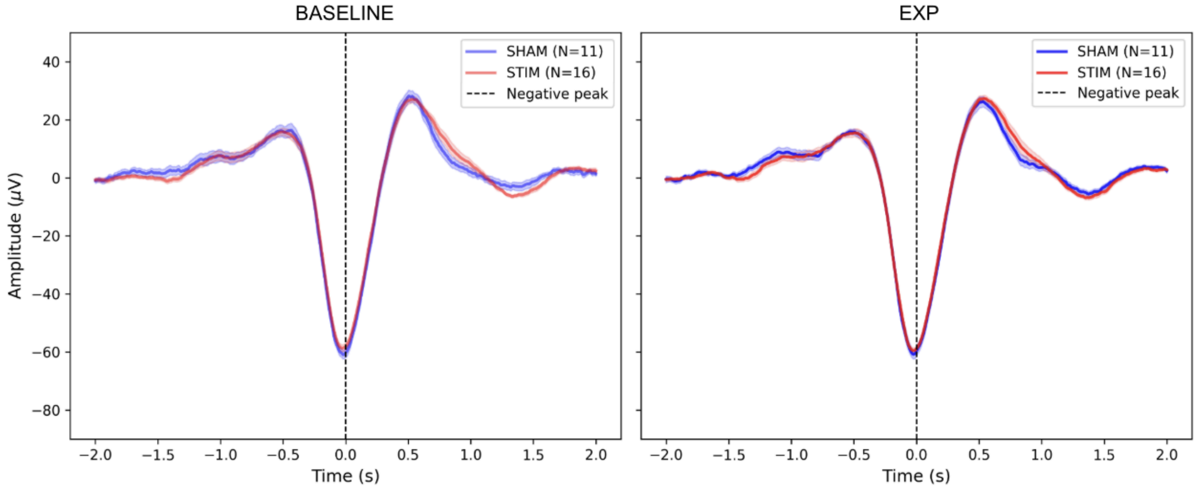

**Figure S6. Average global slow-wave morphology across groups and nights.** Grand-average waveforms of the offline-detected slow waves centered on the negative peak (0 s) are shown for the SHAM (blue, N = 11) and STIM (red, N = 16) groups during the baseline (left) and experimental (EXP, right) nights. Average waveforms were first computed at the subject level after randomly subsampling SOs to match the number of events between BASELINE and EXP within each subject, and group averages were then obtained across subjects.

**Global sleep analyses**

To characterize global sleep patterns, we analyzed sleep architecture across the BASELINE and EXP nights in the SHAM and STIM groups (**Table S6**). Mixed repeated-measures ANOVAs revealed no significant group x night interactions for total sleep time (TST), N1, N2, N3 or NREM sleep duration, wake after sleep onset (WASO), sleep latencies, awakenings, or sleep fragmentation index (SFI).

Sleep latency showed a significant main effect of night ($F(1, 25) = 8.17$, $p = 0.008$), with shorter latencies during EXP in both groups. REM sleep duration showed a significant group x night interaction ($F(1, 25) = 5.81$, $p = 0.024$). REM duration increased by 22.50 ± 6.91 min in SHAM and by 0.66 ± 5.82 min in STIM. Simple-effects analyses indicated a significant increase from BASELINE to EXP within SHAM ($F(1, 10) = 10.60$, $p = 0.009$), whereas the corresponding night effect within STIM was not significant ($p > 0.75$). Between-group differences in REM duration were not significant during either night (both $p > 0.05$).

Compositional analysis did not provide evidence of a between-group difference in the change in the relative proportions of WASO, N1, NREM, and REM (ILR change-score MANOVA: Pillai's trace = 0.197, $F(3, 23) = 1.877$, $p = 0.162$). Similarly, the beta mixed-effects model showed no significant group x night interaction for the proportion of N3 within NREM sleep (b = -0.127, z = -0.366, p = 0.714), providing no evidence of differential change in the relative allocation of NREM sleep to N2 and N3.

**Table S6. Sleep architecture.**

| Variable | ΔSHAM | ΔSTIM | Night | Group | Interaction |
|---|---|---|---|---|---|
| **TST (min)** | 7.32 ± 25.12 | -6.38 ± 18.25 | F(1,25)=0.00, p=0.958 | F(1,25)=0.04, p=0.851 | F(1,25)=0.20, p=0.655 |
| **N1 (min)** | -6.59 ± 5.65 | 0.38 ± 3.30 | F(1,25)=0.67, p=0.422 | F(1,25)=0.17, p=0.680 | F(1,25)=1.29, p=0.267 |
| **N2 (min)** | 11.55 ± 12.12 | -3.16 ± 10.97 | F(1,25)=0.12, p=0.732 | F(1,25)=0.09, p=0.763 | F(1,25)=0.78, p=0.385 |
| **N3 (min)** | 12.05 ± 7.01 | 0.28 ± 3.73 | F(1,25)=1.99, p=0.171 | F(1,25)=0.01, p=0.941 | F(1,25)=2.58, p=0.121 |
| **REM (min)** | 22.50 ± 6.91 | 0.66 ± 5.82 | F(1,25)=4.60, **p=0.042** | F(1,25)=0.01, p=0.912 | F(1,25)=5.81, **p=0.024** |
| **NREM (min)** | 23.59 ± 16.11 | -2.88 ± 11.34 | F(1,25)=0.71, p=0.407 | F(1,25)=0.09, p=0.771 | F(1,25)=1.92, p=0.178 |
| **WASO (min)** | -30.27 ± 15.03 | -3.25 ± 10.09 | F(1,25)=2.78, p=0.108 | F(1,25)=0.02, p=0.894 | F(1,25)=2.41, p=0.133 |
| **Sleep latency (min)** | -13.36 ± 7.11 | -9.06 ± 4.13 | F(1,25)=8.17, **p=0.008** | F(1,25)=0.01, p=0.918 | F(1,25)=0.31, p=0.581 |
| **N3 latency (min)** | -30.91 ± 26.66 | -5.94 ± 9.19 | F(1,25)=1.78, p=0.194 | F(1,25)=0.05, p=0.824 | F(1,25)=1.03, p=0.319 |
| **REM latency (min)** | -5.59 ± 18.28 | 9.84 ± 12.61 | F(1,25)=0.11, p=0.739 | F(1,25)=0.06, p=0.810 | F(1,25)=0.52, p=0.478 |
| **Awakenings** | -3.45 ± 3.37 | 1.69 ± 2.57 | F(1,25)=0.04, p=0.844 | F(1,25)=1.84, p=0.188 | F(1,25)=1.52, p=0.229 |
| **SFI** | -0.89 ± 1.16 | 0.85 ± 0.72 | F(1,25)=0.05, p=0.824 | F(1,25)=1.85, p=0.186 | F(1,25)=1.82, p=0.189 |
| **Variable** | **ΔSHAM** | **ΔSTIM** | **Statistic** | | |
| **N1 (%)** | -2.04 ± 1.30 | -0.13 ± 0.86 | Pillai’s trace=0.197, F(3,23)=1.877, p=0.162 | | |
| **NREM (%)** | 5.49 ± 3.32 | 0.97 ± 1.89 | | | |
| **REM (%)** | 6.24 ± 1.87 | -0.54 ± 1.73 | | | |
| **WASO (%)** | -9.19 ± 4.23 | -0.03 ± 2.68 | | | |
| **Variable** | **ΔSHAM** | **ΔSTIM** | **Group STIM** | **Night EXP** | **Interaction** |
| **N2/NREM (%)** | -4.50 ± 3.24 | -2.40 ± 2.16 | b=0.332, z=0.664, p=0.507 | b=0.259, z=0.927, p=0.354 | b=-0.127, z=-0.366, p=0.714 |
| **N3/NREM (%)** | 4.50 ± 3.24 | 2.40 ± 2.16 | | | |

*Values are expressed as the mean ± SEM of the change between the experimental and baseline nights (Δ = EXP - BASELINE) for the SHAM (N = 11) and STIM (N = 16) groups. Total sleep time (TST), sleep-stage durations, wake after sleep onset (WASO), sleep latencies, awakenings, and sleep fragmentation index (SFI) were analyzed using two-way mixed repeated-measures ANOVAs, with night (BASELINE, EXP) as the within-subject factor and group (SHAM, STIM) as the between-subject factor. Results are reported for the main effects of night and group and their interaction. Between-group differences in changes in the joint composition of wakefulness (W), N1, NREM, and REM were assessed using MANOVA of within-subject changes in isometric log-ratio (ILR) coordinates, with Pillai’s trace reported. The proportion of N3 within NREM was analyzed using a beta mixed-effects model with a subject-specific random intercept. Regression coefficients (b), Wald z-statistics, and p-values are reported for group, night, and their interaction. Coefficients are expressed on the logit scale. SHAM and BASELINE were the reference categories: the group coefficient represents the STIM - SHAM difference during BASELINE, the night coefficient represents the EXP - BASELINE change within SHAM, and the interaction represents the between-group difference in that change. Because N2/NREM and N3/NREM are complementary proportions, only N3/NREM was modeled. NREM denotes N2 + N3. Significant p-values are highlighted in bold.*

To further characterize global sleep physiology, we analyzed relative spectral power across sleep stages and frequency bands. Relative power was calculated as the power within each frequency band normalized by the total spectral power. Mixed repeated-measures ANOVAs revealed no significant main effects of group or night, nor any group x night interactions, for any frequency band or sleep stage (**Table S7**).

**Table S7. Relative spectral power.**

| Sleep stage | Frequency band | ΔSHAM | ΔSTIM | Night | Group | Interaction |
|---|---|---|---|---|---|---|
| NREM | SO | 0.00 ± 0.02 | 0.01 ± 0.01 | F(1,25)=0.60, p=0.444 | F(1,25)=0.00, p=0.980 | F(1,25)=0.37, p=0.548 |
| | Delta | -0.00 ± 0.01 | -0.01 ± 0.01 | F(1,25)=0.38, p=0.544 | F(1,25)=0.51, p=0.484 | F(1,25)=0.07, p=0.797 |
| | Theta | 0.00 ± 0.00 | -0.01 ± 0.00 | F(1,25)=0.49, p=0.490 | F(1,25)=0.62, p=0.437 | F(1,25)=1.20, p=0.283 |
| | Slow spindles | 0.00 ± 0.00 | -0.00 ± 0.00 | F(1,25)=0.01, p=0.937 | F(1,25)=0.03, p=0.869 | F(1,25)=1.60, p=0.218 |
| | Fast spindles | -0.00 ± 0.00 | -0.00 ± 0.00 | F(1,25)=0.10, p=0.756 | F(1,25)=1.18, p=0.287 | F(1,25)=0.13, p=0.720 |
| REM | Theta | 0.01 ± 0.01 | 0.00 ± 0.01 | F(1,25)=0.46, p=0.505 | F(1,25)=1.43, p=0.243 | F(1,25)=0.27, p=0.610 |
| | Beta | 0.01 ± 0.01 | 0.01 ± 0.01 | F(1,25)=3.24, p=0.084 | F(1,25)=0.13, p=0.723 | F(1,25)=0.04, p=0.852 |

*Values are expressed as the mean ± SEM of the change between the experimental and baseline nights (Δ = EXP - BASELINE) for the SHAM (N = 11) and STIM (N = 16) groups. Statistical analyses were performed using two-way mixed repeated-measures ANOVAs with night (BASELINE, EXP) as the within-subject factor and group (SHAM, STIM) as the between-subject factor. P values for the main effects of night and group and for the group x night interaction are shown.*

**Automatic Sleep Staging Performance**

Automatic sleep staging performance was evaluated using 11 experimental-night recordings from the SHAM group for which expert manual scorings were available. Manual sleep staging was performed according to the Rechtschaffen and Kales (R&K) criteria, whereas automatic staging was obtained using the YASA sleep staging algorithm following AASM criteria. Therefore, to enable direct comparison between both scoring systems, stages were mapped as follows: Wake (W) + Movement Time (MT) ↔ Wake (W), Stage 1 (S1) ↔ N1, Stage 2 (S2) ↔ N2, Slow-Wave Sleep (SWS; S3+S4) ↔ N3, REM ↔ REM. Epochs containing at least 10 s of interpolated samples that were classified as artifacts (ART) in the automatic scoring were excluded from the analysis. All epochs labeled as ART corresponded to either W or MT in the manual scorings (**Fig S7**).

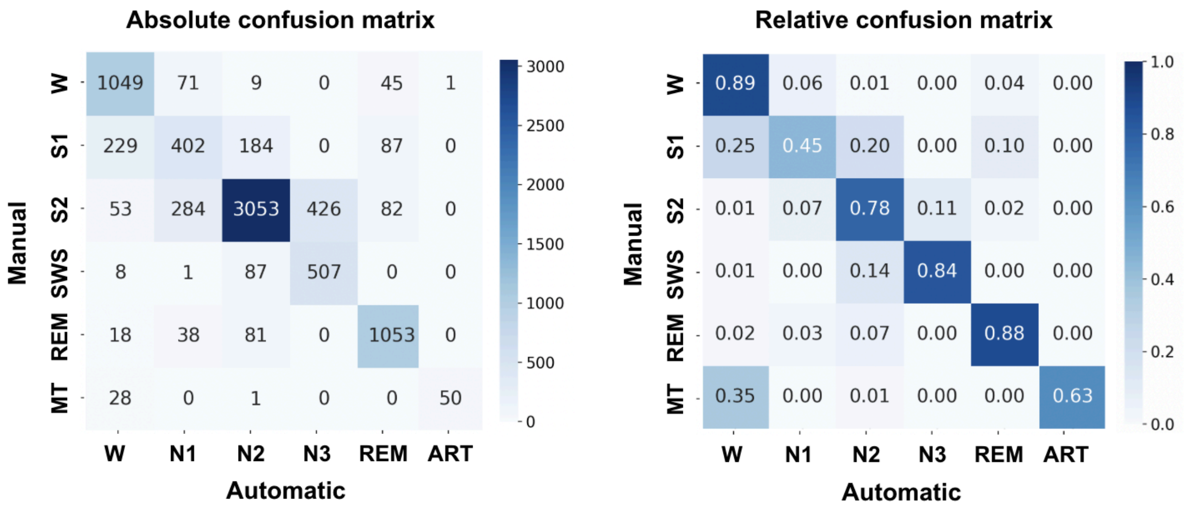

**Figure S7. Confusion matrix comparing raw automatic sleep staging (YASA) and expert manual sleep scoring.** Left: absolute number of 30-s epochs assigned to each stage. Right: row-normalized confusion matrix showing the proportion of epochs assigned to each automatic stage for a given manually scored stage.

Performance was evaluated under two different classification schemes:

- Five-stage comparison: W+MT, S1, S2, SWS, REM versus W, N1, N2, N3, REM.
- Three-stage comparison: W+MT+S1, S2+SWS, REM versus W+N1, N2+N3, REM.

Relative confusion matrices were computed to compensate for class imbalance, particularly the predominance of N2/S2 sleep and the lower representation of SWS/N3 epochs. For each classification scheme, precision, recall, F1-score and overall accuracy were calculated.

*Five-stage classification performance*

**Figure S8** shows the confusion matrix obtained for the five-stage comparison. Overall classification accuracy reached 0.78 across 7,796 analyzed epochs. Agreement was highest for Wake, N2, N3 and REM sleep, while N1 exhibited substantially lower performance. Specifically, F1-scores were 0.83 for Wake, 0.47 for N1, 0.83 for N2, 0.66 for N3, and 0.86 for REM sleep (**Table S8**).

Most classification errors occurred at stage-transition boundaries. N1 epochs were frequently classified as Wake (0.25) or N2 (0.20), whereas N3 epochs were occasionally classified as N2 (0.14). These error patterns closely resemble those reported in the original YASA validation study and are consistent with both the intrinsically low inter-rater agreement of N1 sleep staging and the age-related reduction in slow-wave amplitude, which often results in borderline N2/N3 epochs that are difficult to classify consistently even among expert scorers.

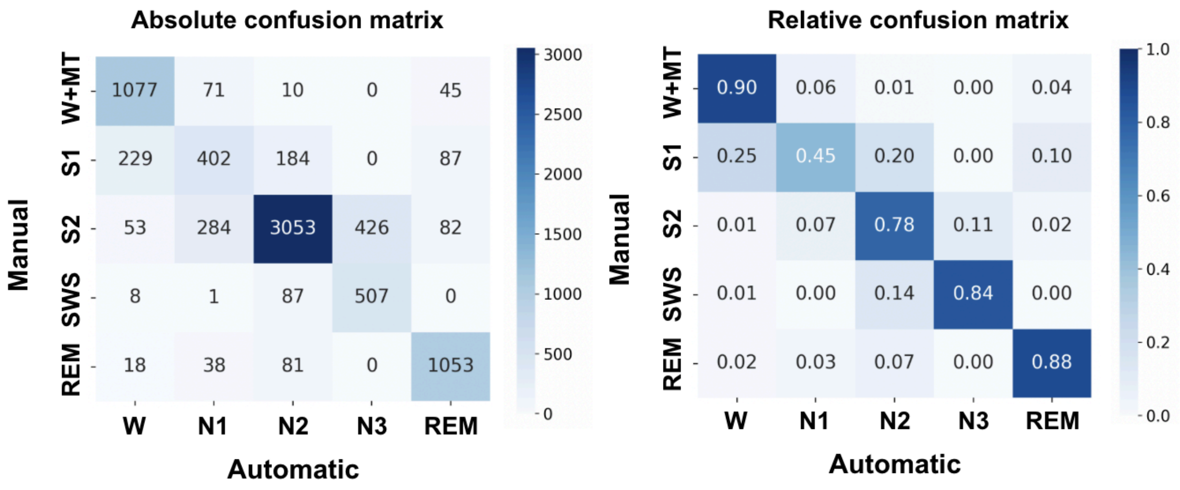

**Figure S8. Confusion matrix comparing automatic sleep staging (YASA) and expert manual sleep scoring using a five-stage classification scheme**. Left: absolute number of 30-s epochs assigned to each stage. Right: row-normalized confusion matrix showing the proportion of epochs assigned to each automatic stage for a given manually scored stage.

**Table S8. Performance metrics for the five-stage sleep staging comparison.**

| Stage | Precision | Recall | F1-score | Support |
|---|---|---|---|---|
| **W** | 0.78 | 0.90 | 0.83 | 1203 |
| **N1** | 0.51 | 0.45 | 0.47 | 902 |
| **N2** | 0.89 | 0.78 | 0.83 | 3898 |
| **N3** | 0.54 | 0.84 | 0.66 | 603 |
| **REM** | 0.83 | 0.88 | 0.86 | 1190 |

*Three-stage classification performance*

To evaluate performance under the broader sleep-stage groups more relevant for the analyses of the present study, stages were further combined into Wake/Light Sleep, NREM Sleep, and REM Sleep. **Figure S9** shows the resulting confusion matrix. Under this classification scheme, overall accuracy increased to 0.89. F1-scores reached 0.83 for Wake/Light Sleep, 0.92 for NREM Sleep, and 0.86 for REM Sleep (**Table S9**). The NREM category showed particularly high agreement with manual scoring. These results indicate that the automatic staging procedure provides robust identification of consolidated NREM sleep periods, which constitute the primary focus of the present analyses.

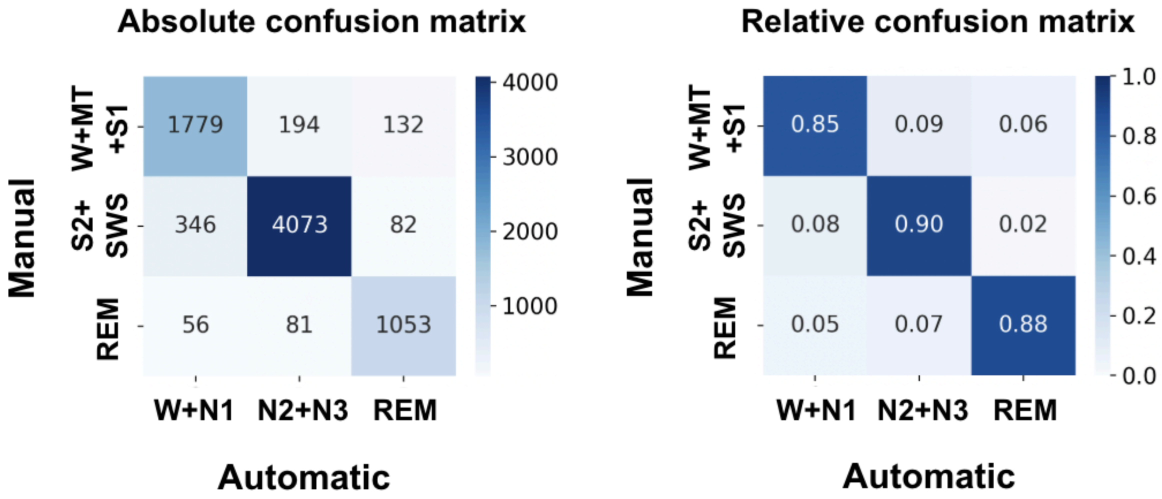

**Figure S9. Confusion matrix comparing automatic sleep staging (YASA) and expert manual sleep scoring using a three-stage classification scheme.** Left: absolute number of 30-s epochs assigned to each stage. Right: row-normalized confusion matrix.

**Table S9. Performance metrics for the three-stage sleep staging comparison.**

| Stage | Precision | Recall | F1-score | Support |
|---|---|---|---|---|
| **Wake/Light Sleep** | 0.82 | 0.85 | 0.83 | 2105 |
| **NREM Sleep** | 0.94 | 0.90 | 0.92 | 4501 |
| **REM Sleep** | 0.83 | 0.88 | 0.86 | 1190 |
